# Implicit Neural Representation of Multi-shell Constrained Spherical Deconvolution for Continuous Modeling of Diffusion MRI

**DOI:** 10.1101/2024.08.30.609148

**Authors:** Tom Hendriks, Anna Vilanova, Maxime Chamberland

## Abstract

Diffusion magnetic resonance imaging (dMRI) provides insight into the micro and macro-structure of the brain. Multi-shell multi-tissue constrained spherical deconvolution (MSMT-CSD) models the underlying local fiber orientation distributions (FODs) using the dMRI signal. While generally producing high-quality FODs, MSMT-CSD is a voxel-wise method that can be impacted by noise and produce erroneous FODs. Local models also do not use the spatial correlation between neighboring voxels to increase parameter estimating power. Additionally, voxel-wise methods require interpolation at arbitrary locations outside of voxel centers. These interpolations can be computationally costly or inaccurate, depending on the method of choice. Expanding upon previous work, we apply the implicit neural representation (INR) methodology to the MSMT-CSD model. This results in an unsupervised machine learning framework that generates a continuous representation of a given dMRI dataset. The input of the INR consists of coordinates in the volume, which produce the spherical harmonics coefficients parameterizing an FOD at any desired location. A key characteristic of our model is its ability to leverage spatial correlations in the volume, which acts as a form of regularization. We evaluate the output FODs quantitatively and qualitatively in synthetic and real dMRI datasets and compare them to existing methods.

## 1 Introduction

Diffusion-weighted magnetic resonance imaging (dMRI) uses directional magnetic gradients to measure the diffusion of water molecules in that direction (Novikov et al., 2019). These acquisitions are sensitive to diffusion at the micrometer scale. Therefore, one can gain insight into the (micro-)structure of the underlying tissue by combining many of these directional acquisitions In dMRI, a b-value relates to the degree of diffusion weighting applied as a function of amplitude (G), time (*δ*), and duration between the gradients (Δ). Using multiple b-values, or shells, provides additional information, as different tissue types respond differently to increasing or decreasing b-value (Genc et al., 2020; Schilling et al., 2017). The dMRI signal can be used to estimate the parameters of an ever-increasing selection of local (voxel-level) microstructural models (Alexander et al., 2019). Each model provides different insights and has its strengths and weaknesses. For white matter, it is particularly interesting to know how fiber bundles are oriented within a voxel. If there are multiple bundles, which is estimated to occur in 60 to 90 percent of voxels (Jeurissen et al., 2013), it is valuable to know how these bundles are angled and what their relative sizes are (Jeurissen et al., 2013). Once known, the fiber orientations can be used to evaluate the local properties directly or in downstream tasks such as tractography. Tractography aims to reconstruct macro-level fiber bundles by computing streamlines that follow the local fiber orientation (Jeurissen et al., 2019).

Constrained spherical deconvolution provides a method to calculate the fiber orientation distributions (FODs) for every individual voxel in a single-shell acquisition (J.-D. Tournier et al., 2007). This method was later expanded to multi-shell multi-tissue CSD (MSMT-CSD), which uses multi-shell data to parameterize FODs for multiple tissue types (Jeurissen et al., 2014). Both methods require the estimation of response functions, which represent the expected signal for different tissues and b-values. CSD only requires a WM response function (obtained from voxels with a single fiber population), while MSMT-CSD requires a response function for every tissue and b-value. The response functions are deconvolved from the dMRI signal to obtain FODs. These FODs are most commonly used for tractography algorithms to reconstruct the structural architecture of the brain or to make inferences about the local microstructure. Using (MSMT-)CSD often results in high-quality (i.e., sharp) FODs, an accurate estimation of partial volume effects, which will positively impact the results of tractography (Jeurissen et al., 2014). Therefore, this technique has become a staple in analyzing dMRI data. However, (MSMT-)CSD operates at the voxel level without using the structural continuity of the brain tissue. This means it is susceptible to noise, as a single noisy voxel can be completely different from and misaligned with its neighbors. This can negatively affect downstream tasks (e.g., fiber tracking, Maier-Hein et al., 2017). Fitting the FODs at locations where data quality is not optimal by using the information in the neighboring voxels (i.e., in noisy environments or low angular and spatial resolution) could help address this issue. Finally, for tractography, it is necessary to calculate FODs in locations that are not voxel center points. In the case of (MSMT-)CSD, these are not readily available, requiring interpolating the dMRI signal or the FODs. Interpolation is computationally costly, and interpolating FODs might produce artificial peaks in areas with strongly curved fibers (Nie & Shi, 2023).

Recently, many machine learning models have been proposed to increase the quality of the fit of the local model (Karimi, 2024). For FOD estimation, most approaches are based on *supervised learning*, which requires a training dataset to learn the model parameters. While promising, these methods are not always ideal: they often require a large amount of data that might impose unwanted biases on the results upon inference, leading to erroneous predictions. To circumvent this problem, *unsupervised* methods, such as Elaldi et al., 2021 and Elaldi et al., 2024, are fit on a single dMRI dataset. Using a UNet-type architecture, the model has access to the spatial information of the dMRI signal to create accurate FOD estimations. However, this approach does not continuously represent the underlying volume as UNets operate in the discretized voxel space. These models, therefore, do not solve the interpolation problems. Implicit neural representations (INRs) provide a way to use neural networks to represent a dataset continuously. Successful usage of INRs outside of the field of dMRI is seen in neural radiance fields, which capture 3D scenes using sparsely sampled 2D images as input (Mildenhall et al., 2020). The robustness to noise and accurate reconstruction of unseen parts of the scene makes INRs a promising avenue of research in dMRI, which deals with noisy, sparsely sampled data. INRs have already been used to compress dMRI data (Mancini et al., 2022), to model dMRI signal (Hendriks et al., 2023), and to model diffusion ODFs (Consagra et al., 2024). While showing promising results, Hendriks et al., 2023 has the limitation of modeling the diffusion signal directly, which limits the usability of the model output as further modeling is required. The diffusion ODFs output by the model from Consagra et al., 2024 are more informative but also require post-processing (i.e., sharpening) to be used in fiber tracking. Additionally, the approach is limited to using single-shell data.

In our work, we propose a method that benefits from the strengths of the CSD and the INR frameworks. We first introduce the methodology for creating a continuous parameterization of CSD and MSMT-CSD using INRs. This results in an unsupervised, response function-informed machine learning model representing a single (multi-shell) dMRI dataset. This continuous model can be sampled at any coordinate to obtain an FOD directly usable for fiber tracking or microstructural analysis. We evaluate the correctness of the learned continuous representations and the robustness of the model to sparsity in both spatial and angular domains, noise, and a changing environment such as different acquisitions, scanners, and patients. Finally, we apply our model to a ‘real-world’ quality dataset to show its application in a realistic scenario.

## 2 Methods

In the upcoming sections, we first briefly overview the (MSMT-)CSD model (section 2.1). Then, we expand upon previous work using INRs to create a novel approach that fits the parameters of the (MSMT-CSD) model in a continuous volume and can leverage the spatial correlations for parameter estimation (section 2.2). The goal is to create an INR that outputs FODs ready for downstream analyses (i.e., to extract metrics (Raffelt et al., 2012) or perform tractography) at any desired location in the volume.

### 2.1 Constrained Spherical Deconvolution

CSD estimates the distribution of fiber orientations present within each voxel. The distribution of these fiber orientations can be modeled with an FOD, a spherical function with an amplitude correlating to the density of the fibers in a given direction (*θ, ϕ*).

In diffusion MRI, these spherical functions are typically described using a truncated, real-valued, even ordered, spherical harmonics (SH) series. We use a basis identical to the one used in MRtrix3 (J.-D. Tournier et al., 2019):

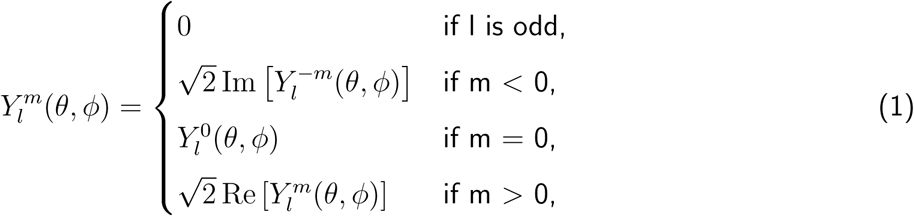

where 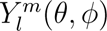 is the SH basis function of order *l* and phase^1^ *m*. Any spherical function can then be represented as:

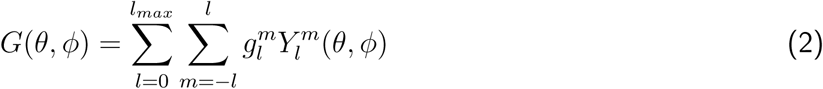

where *G*(*θ, ϕ*) is a generic spherical function, 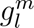 is the SH coefficient for the SH basis function with order *l* and phase *m*, and *l_max_* is the maximum order used in the SH series. The number of coefficients necessary to parameterize the SH-series depends on *l_max_* and can be found using 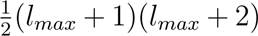.

Given the dMRI signal of a single fiber population, called the ‘response function,’ the dMRI signal in a voxel can be reconstructed by doing a spherical convolution of the single fiber signal with the FOD:

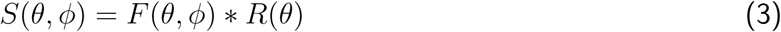

where *S*(*θ, ϕ*) is the measured signal, *F* (*θ, ϕ*) the FOD, and *R*(*θ*) the response function. The estimate for *R*(*θ*) can be obtained from the dMRI data using various existing algorithms. Since *R*(*θ*) is axis-aligned and symmetrical by construction, the spherical harmonic series for *R*(*θ*) only depends on the harmonic phase *m* = 0 terms of the series, the so-called zonal harmonics. The spherical convolution of a spherical harmonics series with a zonal harmonics series is reduced to a simple multiplication of coefficients:

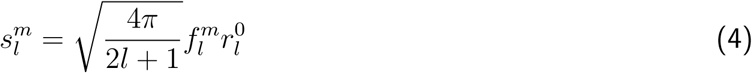

where 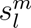 and 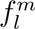 are the SH-coefficients parameterizing *S*(*θ, ϕ*) and *F* (*θ, ϕ*), respectively, 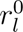 are the zonal harmonics coefficients parameterizing *R*(*θ*). The scaling factor in front corresponds to the l-th order coefficients for the spherical harmonics series for a Dirac-delta function. The FOD can be retrieved by deconvolving the measured dMRI signal with the single fiber population signal. The retrieved FOD is constrained to be strictly positive by penalizing negative values during optimization, thereby filtering high-frequency noise that generates these negative values. Details about the CSD algorithm can be found in J.-D. Tournier et al., 2007. MSMT-CSD expands this idea by using data with multiple b-values and response functions to get more accurate estimations of the FOD and separate volume fractions for other tissue types, such as grey matter (GM) and cerebrospinal fluid (CSF). The method then calculates an orientation distribution for every tissue type. These distributions are convolved with their respective response functions. The response functions in the case of MSMT-CSD are not only parmeterized by *θ*, but also by the b-value. The convolution provides the expected signal for each shell. Since the GM and CSF are presumed to be tissues with (restricted) isotropic diffusion, an orientation distribution of a single coefficient suffices per tissue. These coefficients are often seen as a volume fraction of these tissue types. Details about the MSMT-CSD algorithm can be found in Jeurissen et al., 2014.

### 2.2 Continuous modeling of CSD through Implicit Neural Representation

In our work, we represent a full multi-dimensional dMRI dataset using an INR that outputs FODs for any spatially encoded 3D input coordinate (Figure 1). This borrows concepts from Neural Spherical Harmonics, previously used to model the dMRI signal directly (Hendriks et al., 2023). We extend the existing approach by adapting the model architecture to output FODs and introduce the (MSMT-)CSD framework to relate the FODs to the measured signal. This injects additional assumptions (e.g., the response functions) into the INR fitting process. In the upcoming sections, we expand upon the positional encoding (Figure 1a), the adapted model architecture (Figure 1b), and the fitting of the model (Figure 1c).

**Figure 1:**
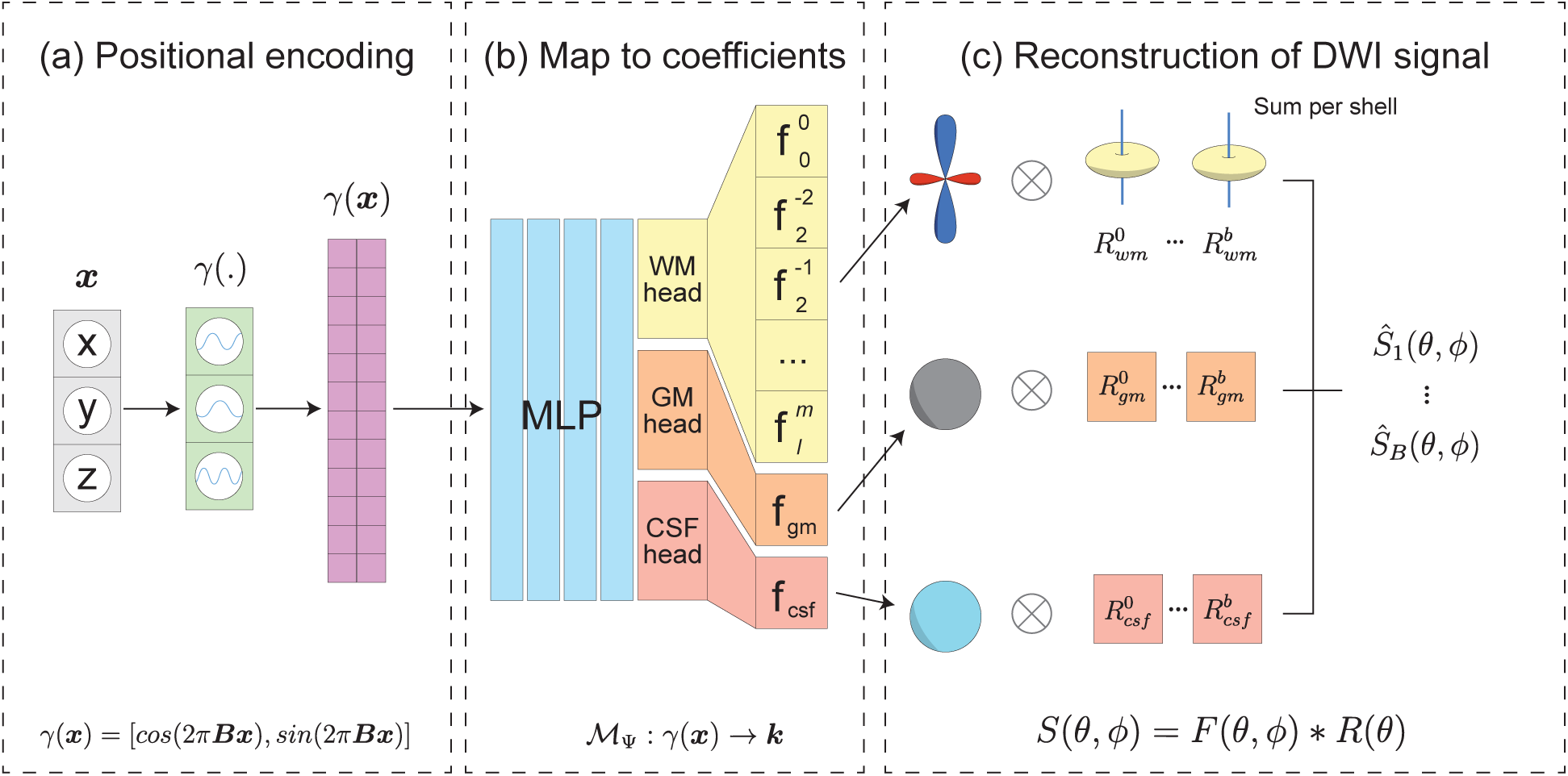
(a) The 3D coordinates are positionally encoded using (co)sine waves of different frequencies randomly sampled from a Gaussian distribution. (b) The architecture of the proposed multi-shell model. In the case of the single shell model, the three output heads are removed, and instead, the coefficients for the FOD are output directly. (c) The signal reconstruction from the different tissue types in the multi-shell model. The output spherical functions (WM FOD, GM, and CSF volume fractions) are convolved with the different response functions for that shell and then summed together to obtain a single output function per shell.

#### Positional encoding

The model input is a 3D-coordinate ***x*** = [*x, y, z*]. Using just the raw coordinates as input leads to a poor representation of the dataset as the model cannot capture the higher frequency details. Previous studies show that encoding the coordinates using a multitude of sine and cosine functions with different frequencies significantly improves the representation (Mildenhall et al., 2020). In this work, we leverage existing work based on frequencies randomly sampled from a Gaussian distribution to prevent an axis alignment bias during encoding, which has shown to be effective at representing different types of datasets (including conventional MRI) (Tancik et al., 2020). The encoding is shown in (5).

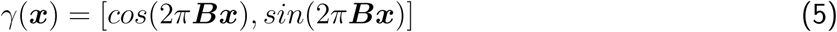

Where *γ*(***x***) returns the encoded coordinate vector ***x*** (which is normalized to lie in range [−1, 1] prior to encoding), and ***B*** is a size *n_p_* × *d* matrix with values sampled from N (0*, σ*^2^). The hyperparameter *n_p_* determines the number of positional encodings, *d* is the number of dimensions in coordinate ***x***, and *σ* is the standard deviation of the Gaussian. The hyperparameters *n_p_* and *σ* can be tuned to generate the desired representation of the data, where *n_p_* is more related to dataset complexity, and *σ* to the granularity of the representation (i.e. higher values for *σ* capture finer details, but are more noisy) (Tancik et al., 2020).

#### Model architecture

We define a multi-layer perceptron (MLP) that maps positionally encoded input coordinate *γ*(***x***) to an SH-coefficient vector ***k*** (6)

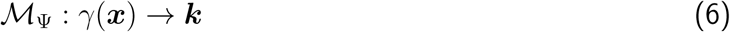

where M_Ψ_ is the MLP parameterized with weights Ψ, and ***k*** is given by:

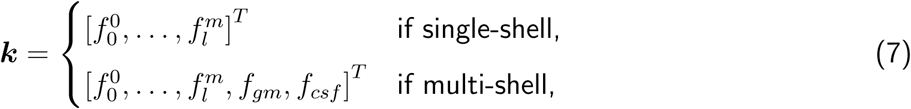

where 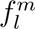 is the vector of coefficient of the SH basis function of order *l* and phase *m*, and *f_gm_* and *f_csf_* are the coefficients for the GM and CSF, respectively.

We use two similar but slightly different model architectures for the single-shell and multi-shell implementation. Both models take *n_p_* positionally-encoded 3D coordinates as an input, followed by an MLP consisting of four layers with ReLU activation functions. The size of each layer is a tunable hyperparameter. For the single-shell model, the MLP directly outputs SH-coefficients that parameterize the FOD at the input coordinate. For the multi-shell model, the output of the MLP is used as input for three output heads, one for each tissue type, consisting of one fully connected layer. These heads are optimized alongside the MLP during model fitting. The WM head outputs a full SH-series parameterizing the FOD, while the GM and CSF heads output a single coefficient. The complete model architecture can be seen in Figure 1b.

#### Model fitting

The model fits the dataset by calculating a mean-squared error (MSE) between the measured diffusion signal *S*(*θ, ϕ*) and the diffusion signal reconstructed from the FODs 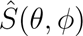. In single-shell CSD, 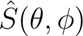 is reconstructed by performing a convolution between the output FOD and the response function as shown in (4). This model will be referred to as INR-CSD.

For MSMT-CSD, the model outputs 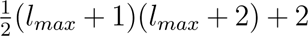 coefficients (a full SH-series for the WM and a zeroeth order SH-series for both GM and CSF). The signal is obtained for each b-value individually as the sum over *T* tissue types (three in the case of MSMT-CSD) of the convolution of the tissue FODs with the corresponding tissue response functions belonging to that b-value. The multi-shell output for a single measured coordinate, therefore, is:

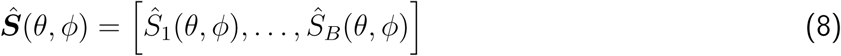

where *B* is the number of b-values and 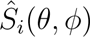 is the signal for b-value *i*. We obtain 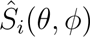 from

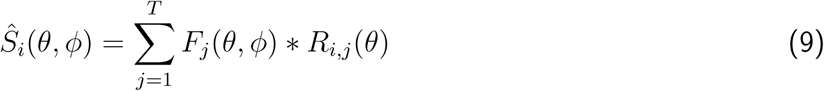

where *F_j_* (*θ, ϕ*) is the FOD for tissue type j, parameterized by the corresponding coefficients in ***k***, and *R_i,j_* (*θ*) is the response function for b-value *i* and tissue type *j*. This process is shown in Figure 1(b, c). This model will be referred to as INR-MSMT.

Since ***k*** and therefore *F_j_*, are dependent on 3D coordinate ***x***, we can define the signal in the entire volume as:

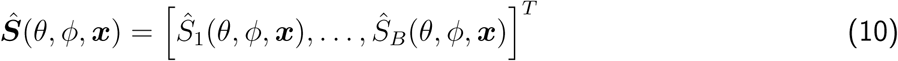

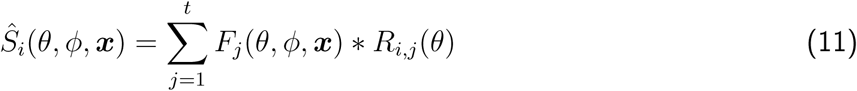

Finally, we constrain the FOD(s) to be strictly positive as described in J.-D. Tournier et al., 2007. The (WM) FOD is sampled in 724 directions, and any values larger than 0.1 times the mean amplitude of the FOD are included in the loss. The GM and CSF are, by construction, non-negative.

Given a set of b-vectors with directions (*θ, ϕ*) in *B* b-values, where b-value *i* contains *Q_i_* b-vectors,

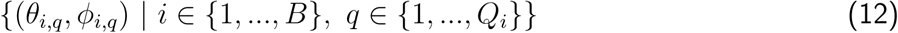

for a set of *N* 3D coordinates ***X*** = {***x*_0_***, …, **x******_N_*** } (one for every voxel in a dMRI dataset), we obtain the parameters for the INR M_Ψ_∗ by minimizing the MSE alongside the constraint:

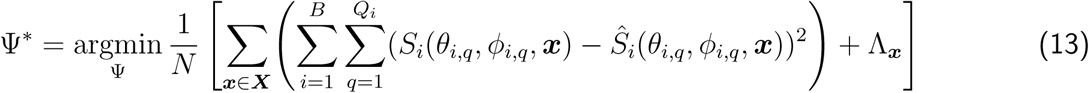

Where 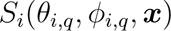 and 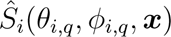 are respectively the measured signal and the reconstructed signal at coordinate ***x*** for b-value *i* in direction (*θ_i,q_, ϕ_i,q_*), and Λ***_x_*** is the non-negativity constraint.

### 2.3 Datasets

We use a synthetic high-resolution brain-like dataset to introduce our methodology and perform a quantitative analysis of its performance regarding sparsity and noise. The diffusion-weighted images are generated using Fiberfox (Neher et al., 2014), from curated tractograms that have been used previously in dataset generation of the 2015 ISMRM challenge (Maier-Hein et al., 2017; Renauld et al., 2023). Since we are interested in both single-shell and multi-shell CSD, the dataset we generate for this experiment has 67 volumes spread over three b-values: 7 for *b* = 0*s/mm*^2^, 30 for *b* = 1200*s/mm*^2^, and 30 for *b* = 3000*s/mm*^2^. For consistency, these are the same gradient directions and b-values used in real brain acquisition in the real-data experiments. We generate the dataset in a 1.25 mm isotropic resolution and add no artifacts or noise. The rest of the settings are identical to those used for generating the challenge data. Additionally, a dataset is generated at 2.5 mm isotropic resolution for the low spatial resolution experiment. Finally, three datasets are generated using Fiberfox at 1.25 mm isotropic resolution with simulated complexGaussian noise, resulting in three datasets with signal-to-noise ratios (SNRs) of 15, 25, and 35 for the noise-related experiments.

Additionally, we use the computational diffusion MRI 2018 (cdMRI2018) challenge preprocessed dataset (Ning et al., 2020; Tax et al., 2019) to explore model performance and robustness in a realistic setting. Two randomly chosen subjects (B and K) were scanned on two scanners (3T Siemens Prisma and Connectom) with two acquisition settings per scanner: standard (ST) and state-of-the-art (SA). This provides a baseline for comparing the performance under varying acquisition parameters. Both scanners have ST acquisitions of 2.4mm isotropic resolution with 67 volumes spread over three b-values: 7 for *b* = 0*s/mm*^2^, 30 for *b* = 1200*s/mm*^2^, and 30 for *b* = 3000*s/mm*^2^. Additionally, the Prisma scanner has a 1.5mm isotropic resolution, 136 volume SA acquisition spread over three b-values: 16 for *b* = 0*s/mm*^2^, 60 for *b* = 1200*s/mm*^2^, and 60 for *b* = 3000*s/mm*^2^. Finally, the Connectom scanner has a 1.2mm isotropic resolution SA acquisition, with volumes identical to the Prisma SA acquisition.

### 2.4 Evaluation

For the experiments using synthetic datasets described below, we perform a quantitative analysis of the resulting FODs by comparing them to the FODs generated on the clean, high-resolution multi-shell dataset using voxel-wise MSMT-CSD. We assume these FODs to be the ground truth (GT). We report the total apparent fiber density (AFD) as a measure of FOD scale, the angular correlation coefficient (ACC) as a measure of angular correctness, and the number of fiber orientations (NuFO) to compare the peaks used for tracking (Anderson, 2005; Dell’Acqua et al., 2013; Raffelt et al., 2012). Furthermore, we visualize the FODs in all experiments to make a qualitative assessment.

#### Implementation

We use *l_max_* = 8 for every SH series in this paper. This choice is informed by previous work showing that higher orders have low contributions to the signal (J.-D. Tournier et al., 2013). The spherical harmonic basis functions are obtained through scipy^2^.

The response functions for single-shell CSD are retrieved from the datasets using the Tournier algorithm, and for the multi-shell algorithm using the Dhollander algorithm, both implemented in MRtrix3 (Dhollander et al., 2019; J.-D. Tournier et al., 2013).

The INR-models in all experiments are implemented in PyTorch^3^. An Adam optimizer was used with a learning rate of 1 × 10^−4^ and default settings otherwise. Further hyperparameter settings are determined in the first experiment and used throughout the rest.

The AFD and NuFO are calculated using scilpy^4^, with an absolute threshold of 0.07 and default settings otherwise. AFD (total) is defined as being the first coefficient of the SH-series (Raffelt et al., 2012).

The ACC is calculated between two SH-series with coefficients 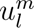 and 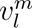 as:

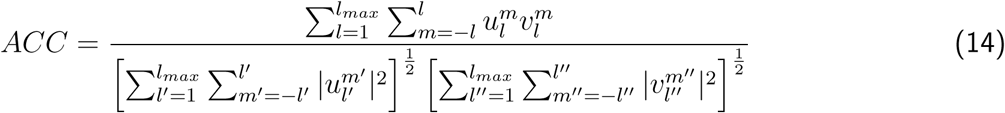

which corresponds with Anderson, 2005 and Schilling et al., 2019. The metrics are computed in all voxels where the ground truth FOD has a first coefficient greater than 0.05 for the WM.

## 3 Experiments

### Implicit neural representation of CSD and MSMT-CSD

The feasibility of our approach is first assessed by fitting the INR-CSD and INR-MSMT models for a full volume. Our approach is feasible if the model can reconstruct the original dMRI signal. The INR-CSD is fit to the *b* = 3000*s/mm*^2^ shell of the 1.25mm isotropic resolution synthetic dataset. The outputs are visualized for an area of interest and compared qualitatively and quantitatively to the GT. This experiment is repeated for INR-MSMT using all three b-values (*b* = 0, 1200, 3000*s/mm*^2^) of the 1.25mm synthetic dataset.

### Low spatial resolution

For the INR to handle spatial sparsity well, it must give valid outputs between the voxel positions for which it was fit. To assess this, we fit an INR-CSD model on the *b* = 3000*s/mm*^2^ and an INR-MSMT on all three b-values of the 2.5mm isotropic synthetic dataset. As a first indication of intervoxel correctness, we visualize the FODs resulting from supersampling the model at 1.25mm isotropic resolution in two regions of interest. These FODs are compared to a set of FODs obtained by linear interpolation of the SH-coefficients of the FODs calculated at 2.5mm resolution using voxel-wise CSD. The AFD and the ACC are computed between both models and the GT and are reported alongside the NuFO. We then explore the intervoxel space by 1) selecting two voxels in each region, 2) sampling the model at multiple equidistant steps between these voxels, and 3) visualizing the FODs at those locations. Additionally, we show maps of the GM and CSF volume fractions for the ground truth, INR-MSMT, and linearly upsampled MSMT-CSD volume fractions.

### Sparsity in angular resolution

To evaluate the performance of the INR in settings with low angular resolution, we iteratively remove b-vectors from the dataset. At every step, we remove the b-vectors with the smallest effect on the hemispherical coverage. Since we expect model failure only to occur at considerably sparse data, the starting point for our evaluation is a single shell of 30 b-vectors at *b* = 3000*s/mm*^2^ in the 1.25mm isotropic synthetic dataset. For every subset of b-vectors, we fit an INR-CSD and a voxel-wise CSD model. The AFD and ACC are calculated between both models and the GT for different numbers of b-vectors. The ACC maps, FODs, and NuFO are also reported for a single set of b-vectors of interest.

### Robustness to noise

We evaluate the robustness of our approach to noise by fitting the INR-CSD to synthetic datasets with an SNR of 15, 25, and 35. The resulting FODs are compared visually to the FODs of voxel-wise CSD. For the dataset with SNR = 25, we report the AFD and the ACC calculated between both models and the GT, and NuFO.

### Robustness of hyperparameter settings

We aim to show the performance of the model across different scanners and acquisitions when using identical hyperparameters. We fit our INR-MSMT model to the cdMRI 2018 dataset (Ning et al., 2020). Two subjects were randomly selected, each with two acquisitions on two scanners, resulting in 8 different INR-MSMT fits. To assess the robustness of our model, we qualitatively analyze the consistency of the FODs within the subject and the difference between subjects by visualizing them. The FODs are compared to FODs obtained through voxel-wise MSMT-CSD.

### Real-world application of the model

Having a multi-shell high-resolution dataset is not common in day-to-day practice. To test the perfomance of the INR on a ‘real-world quality’ dataset, we fit an INR-CSD on a single *b* = 3000*s/m*^2^ shell of the ST of the Connectom scanner and sample it at SA resolution. The resulting FODs are compared to a set of FODs obtained by linear interpolation to SA resolution of the SH-coefficients of the FODs calculated at ST resolution using voxel-wise CSD. We show the MSMT-CSD fit on the SA data as an additional comparison, which corresponds to FODs obtained in a ‘best-case scenario’ in current practice.

## 4 Results

### Implicit neural representation for CSD and MSMT-CSD

The following hyperparameter settings were found using a grid search and used for fitting both models: 1024 neurons per layer, four layers, *σ* = 4, *n_p_* = 5000. The influence of these parameters is detailed in Appendix A. The single-shell and multi-shell INRs converge after approximately 650.000 iterations over 30 epochs with batch size 500 on a 16GB NVIDIA RTX 4080. Training times are 2m 58s and 5m 24s respectively. Figure 2 shows the ACC map and FODs in the centrum semi-ovale (CSO) for the single-shell model alongside the GT FODs. Quantitative results for both models are shown in Table 1. We observe minimal differences between the angles of the fiber populations and the relative sizes for this feasibility experiment, compared to the ground truth. Quantitative analysis confirms that the errors of the INRs in representing the target data are low.

**Figure 2:**
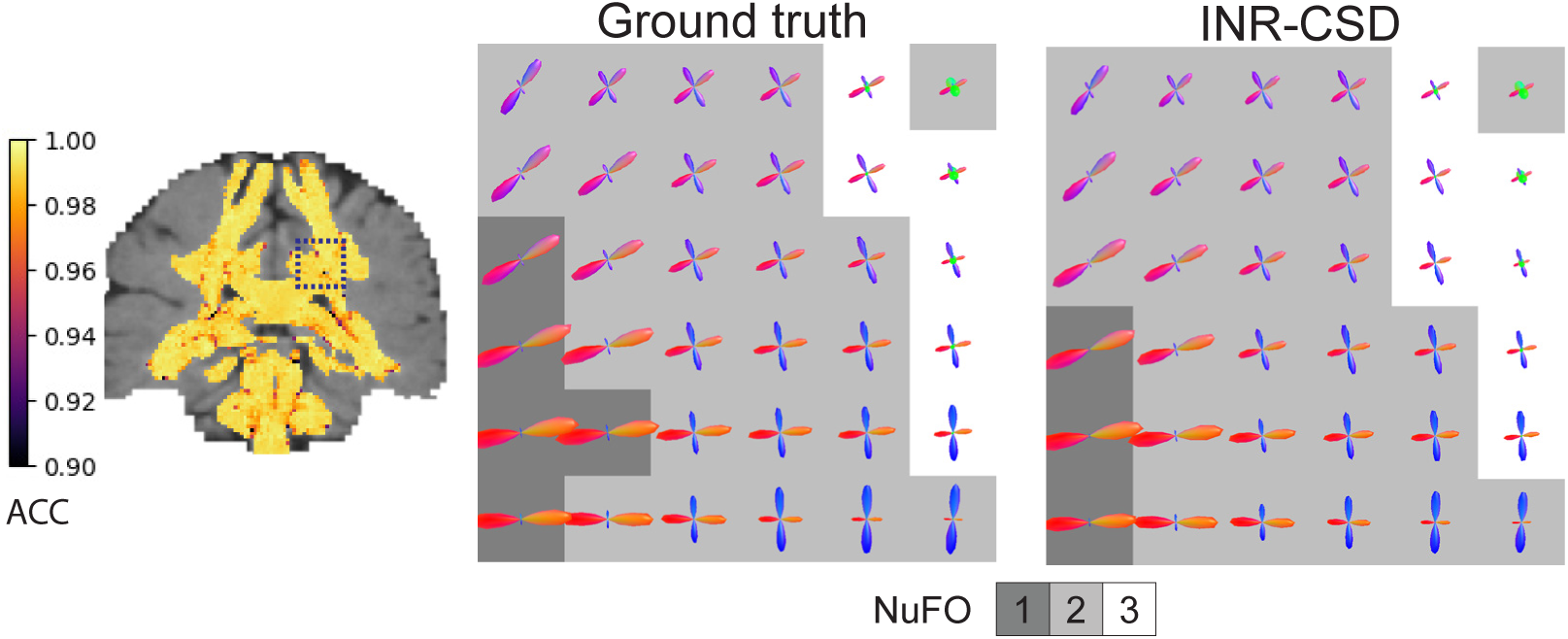
Visualizations of the results for the feasibility experiment on the synthetic dataset. The angular correlation coefficient (ACC) map for INR-CSD is visualized on the left. The FODs are visualized in the centrum semi-ovale (dark blue square in the ACC map) for the ground truth and INR-CSD. The FODs are shown against a background that displays the number of fiber orientations (NuFO) in that voxel.

**Table 1:**
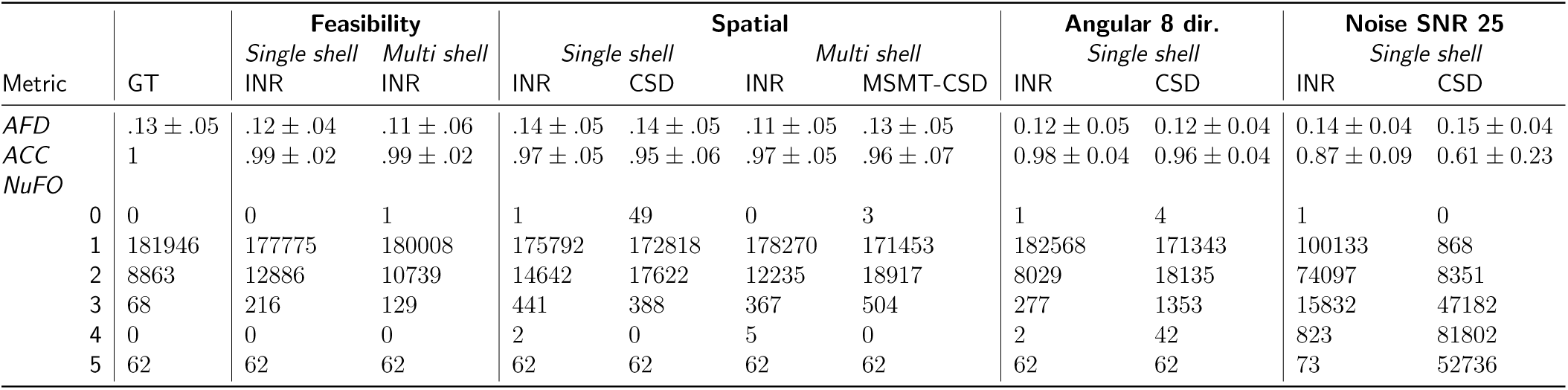
Quantitative results for the synthetic dataset experiments.

| Metric | GT | Feasibility |  | Spatial |  |  |  | Angular 8 dir. |  | Noise SNR 25 |  |
| --- | --- | --- | --- | --- | --- | --- | --- | --- | --- | --- | --- |
|  |  | Single shell<br>INR | Multi shell<br>INR | Single shell<br>INR | CSD | Multi shell<br>INR | MSMT-CSD | Single shell<br>INR | CSD | Single shell<br>INR | CSD |
| AFD | .13 ± .05 | .12 ± .04 | .11 ± .06 | .14 ± .05 | .14 ± .05 | .11 ± .05 | .13 ± .05 | 0.12 ± 0.05 | 0.12 ± 0.04 | 0.14 ± 0.04 | 0.15 ± 0.04 |
| ACC | 1 | .99 ± .02 | .99 ± .02 | .97 ± .05 | .95 ± .06 | .97 ± .05 | .96 ± .07 | 0.98 ± 0.04 | 0.96 ± 0.04 | 0.87 ± 0.09 | 0.61 ± 0.23 |
| NuFO |  |  |  |  |  |  |  |  |  |  |  |
| 0 | 0 | 0 | 1 | 1 | 49 | 0 | 3 | 1 | 4 | 1 | 0 |
| 1 | 181946 | 177775 | 180008 | 175792 | 172818 | 178270 | 171453 | 182568 | 171343 | 100133 | 868 |
| 2 | 8863 | 12886 | 10739 | 14642 | 17622 | 12235 | 18917 | 8029 | 18135 | 74097 | 8351 |
| 3 | 68 | 216 | 129 | 441 | 388 | 367 | 504 | 277 | 1353 | 15832 | 47182 |
| 4 | 0 | 0 | 0 | 2 | 0 | 5 | 0 | 2 | 42 | 823 | 81802 |
| 5 | 62 | 62 | 62 | 62 | 62 | 62 | 62 | 62 | 62 | 73 | 52736 |

### Low spatial resolution

We used hyperparameter settings identical to those used in the previous experiment to fit the INRs. Figure 3(a,b,c) shows the error maps of the AFD for both models alongside the difference in error. Our method has a smaller error in areas where fibers cross but larger in areas with a single fiber population. Figure 3 also shows two regions of interest: an area of crossing fibers and an area of strongly curved fibers. We observe that INR-CSD can correctly interpolate crossing and curving fibers at subvoxel levels. Linear interpolation does not correctly interpolate the FODs in curved fibers, creating spurious peaks as shown in the yellow box in Figure 3. Figure 4 shows maps of the GM and CSF volume fractions of the MSMT-CSD and INR-MSMT fits. Both show a loss of detail compared to the ground truth. Notably, the INR-MSMT has a higher estimate for the GM volume fraction and a lower estimate for the CSF volume fraction in areas where brain tissue is present. The quantitative results in Table 1 show identical AFD values and a slightly larger ACC for the INR-CSD for this low spatial resolution experiment, compared to the ground truth. The INR-MSMT shows a minor underestimation of the AFD, with similar ACC scores. The distribution of the NuFO looks similar across methods, showing an underestimation of single-fiber populations and an overestimation of dual-fiber populations. The overestimation of dual-fiber populations is more pronounced in the interpolated voxel-wise models.

**Figure 3:**
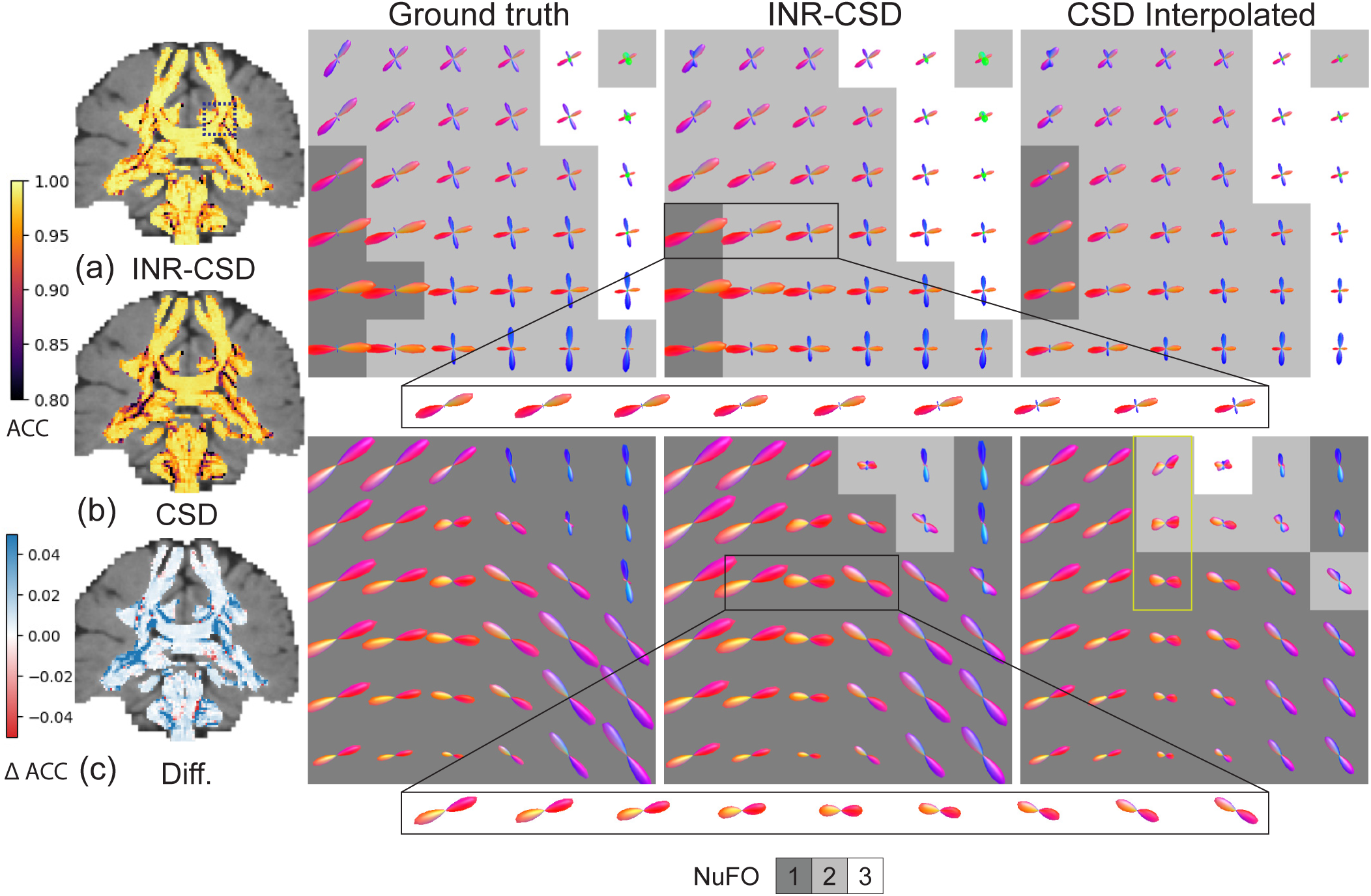
Visualizations of the results for the spatial resolution experiment using the 2.5mm resolution synthetic dataset to fit the models. The angular correlation coefficient (ACC) maps for INR-CSD and CSD (a, b) and their difference (c) are shown. In (c), blue and red indicate a higher ACC in INR-CSD and CSD, respectively. The FODs are visualized in the centrum semi-ovale (CSO, top row, marked in (a) with a blue square) and an area near the ventricle, inferior to the CSO (bottom row) for the ground truth, INR-CSD, and CSD. The FODs are shown against a background that displays the number of fiber orientations (NuFO) in that voxel. The yellow box shows an area where linear interpolation creates spurious peaks.

**Figure 4:**
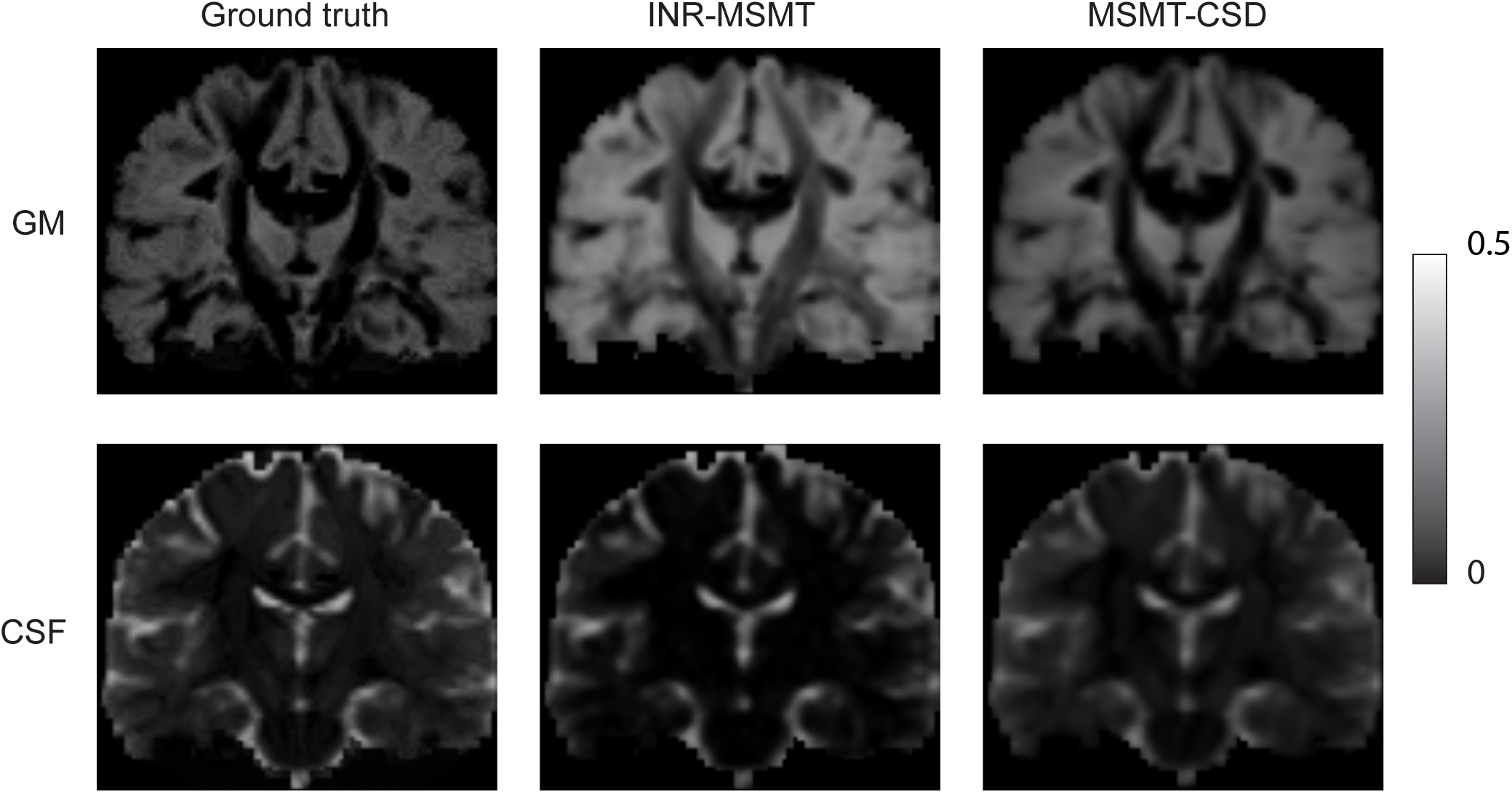
Visualizations of a coronal slice of the grey matter (GM) and cerebrospinal fluid (CSF) volume fraction (*f_gm_*, *f_csf_*) maps for the high-resolution MSMT-CSD (ground truth), the INR-MSMT fit on 2.5mm isotropic resolution and sampled at 1.25mm resolution, and MSMT-CSD fit on 2.5mm resolution linearly interpolated to 1.25mm resolution.

### Sparsity in angular resolution

The mean ACC and the mean average error (MAE) in AFD for both the CSD and INR-CSD model for an increasing number of directions are shown in Figure 5. One can see that the INR-CSD model shows higher ACC and lower MAE in AFD compared to CSD, although differences are marginal. In Figure 6, where the outputs of the models fit on eight directions are shown in an area of low crossing angle (top row), we can see that in regions of low crossing angle, INR-CSD does not find two distinct peaks where CSD (mostly) does (NuFO = 2). Meanwhile, in the area of high curvature (bottom row), we see spurious peaks in the CSD fit, which do not appear in INR-CSD. The ACC maps show an overall higher ACC for the INR-CSD fit, although both models perform sub-optimally in areas of crossing fibers. Table 1 shows an almost identical AFD, while the ACC is marginally higher for INR-CSD for this angular sparsity experiment, compared to the ground truth. NuFO shows a slight shift towards higher numbers for the CSD fit. The first row in Figure 7 shows how spurious peaks show up throughout the WM for CSD.

**Figure 5:**
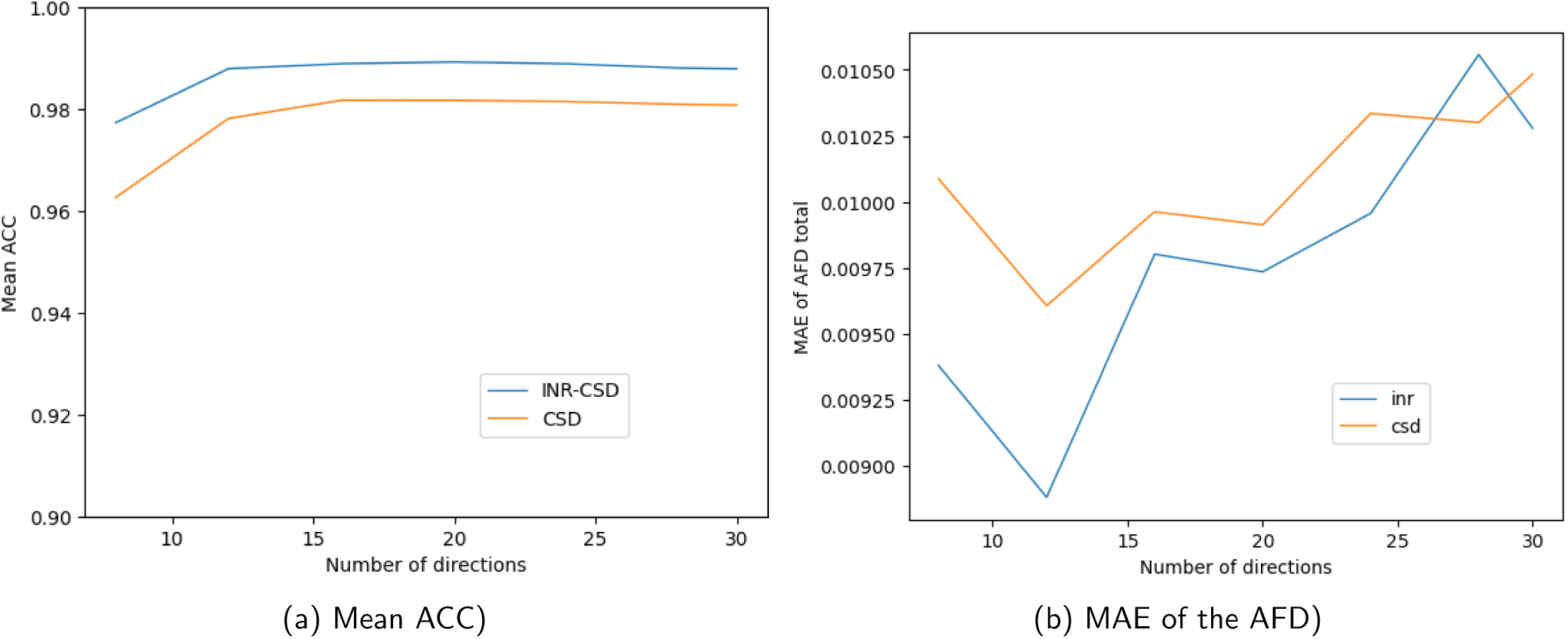
Mean angular correlation coefficient (ACC) and mean average error (MAE) of the apparent fiber density (AFD) for INR-CSD and CSD fit on different numbers of directions of the *b* = 3000*s/m*^2^ shell of the synthetic dataset

**Figure 6:**
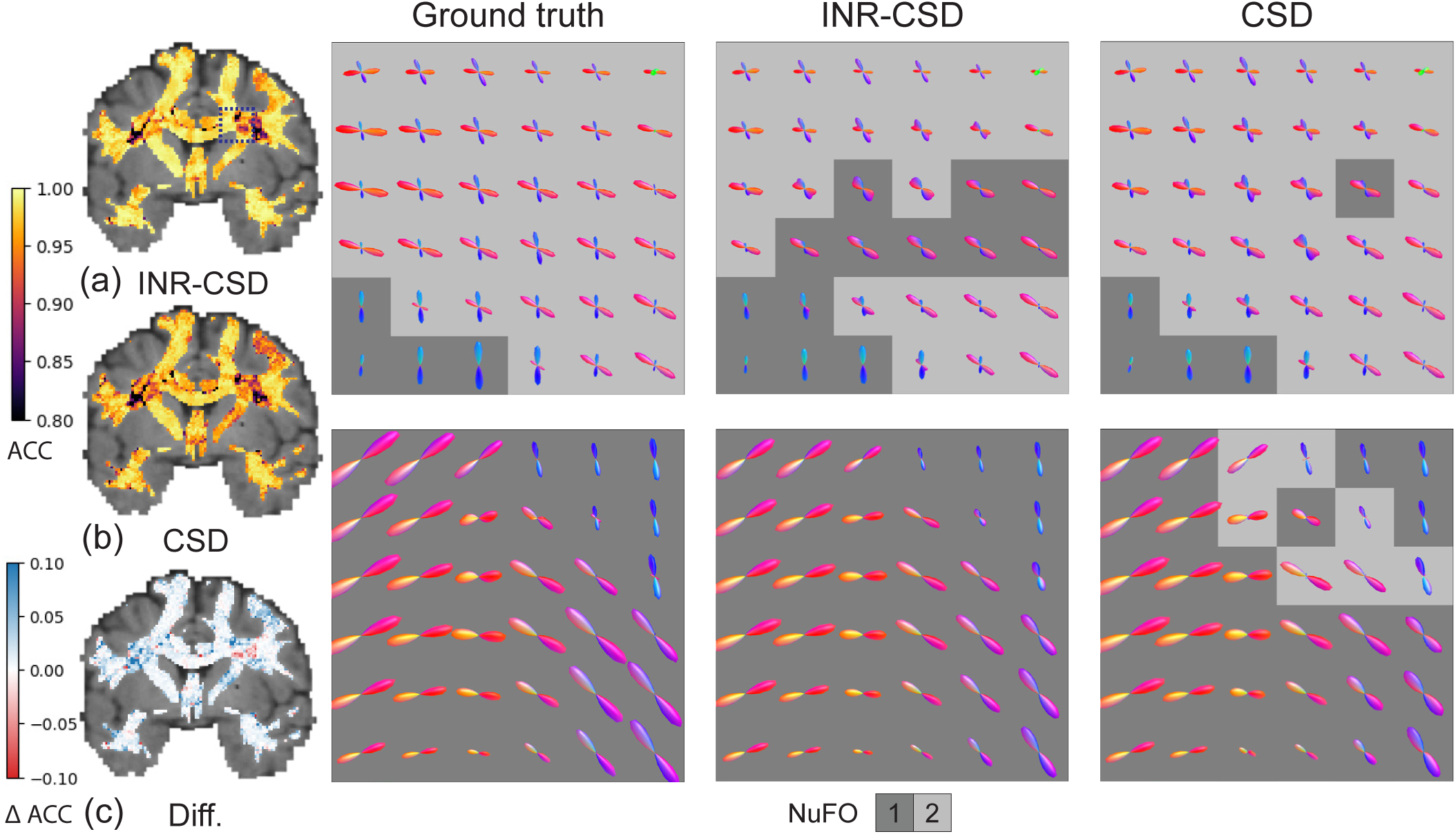
Visualizations of the results for the angular resolution experiment, fitting the models on eight directions of the *b* = 3000*s/m*^2^ shell of the synthetic dataset. The angular correlation coefficient (ACC) maps for INR-CSD and CSD (a, b) and their difference (c) are shown. In (c), blue and red indicate a higher ACC in INR-CSD and CSD, respectively. The FODs are visualized in the centrum semi-ovale (CSO, top row, marked in (a) with a blue square) and an area near the ventricle, inferior to the CSO (bottom row) for the ground truth, INR-CSD, and CSD. The FODs are shown against a background that displays the number of fiber orientations (NuFO) in that voxel.

**Figure 7:**
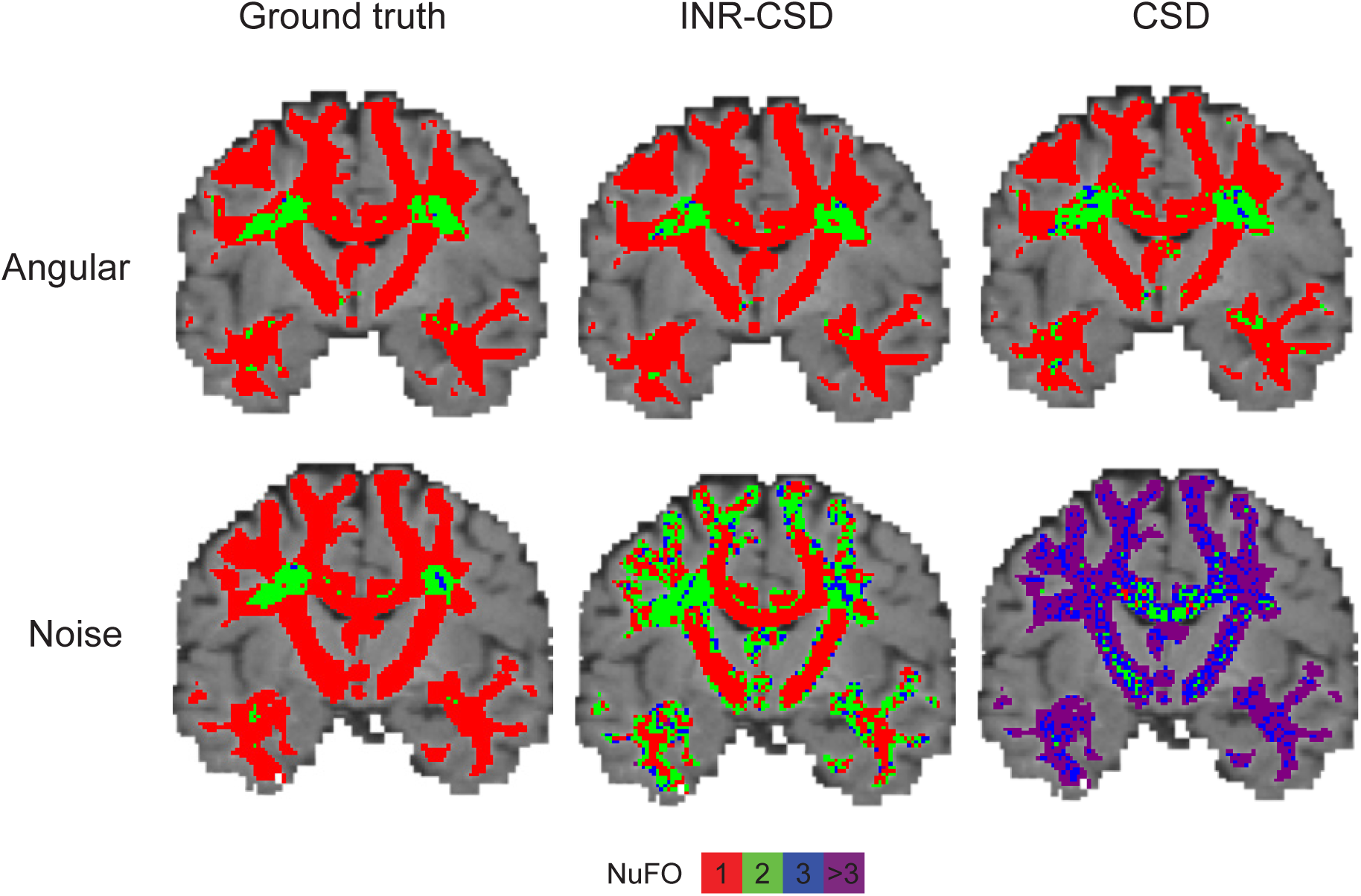
Visualizations of NuFO maps for both the eight directions angular experiment (top row) and SNR = 25 noise experiment (bottom row) for both INR-CSD and CSD and the MSMT-CSD on the full dataset (ground truth).

### Robustness to noise

The FODs in the centrum semi-ovale for different levels of SNR for INR-CSD and CSD are shown in Figure 8. A marked difference can be observed between both methods: the INR-CSD appears to be more robust to noise, showing similar results from SNR = 35 to SNR = 15. In contrast, the FODs obtained through CSD are affected by data quality, especially with lower SNRs. Table 1 shows the quantitative results for SNR = 25, which support the qualitative analysis. The mean ACC for INR-CSD is 0.87, while voxel-wise CSD scores 0.61. NuFO counts show the distribution of fiber orientations of INR-CSD, which more closely resembles the GT but is shifted toward higher population counts. This shift is much more pronounced for the CSD fit, where nearly all voxels have three or more fiber populations. The second row in Figure 7 exemplifies this difference.

**Figure 8:**
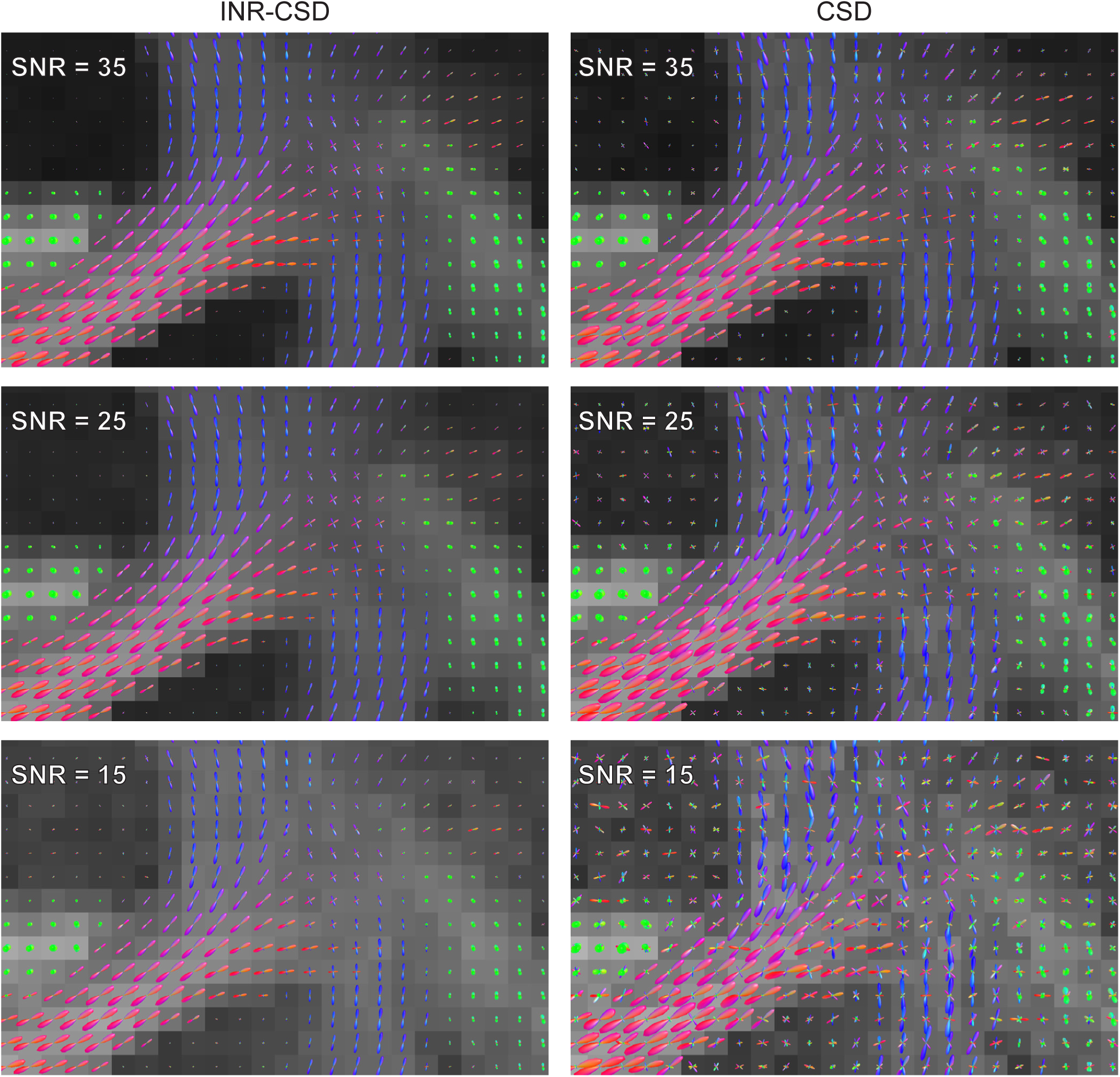
Visualizations of FODs in the centrum semi-ovale for both INR-CSD and CSD fit on the b = 3000s/m2 shell of the synthetic dataset with different signal-to-noise ratios (SNRs), against a background of apparent fiber density.

### Robustness of hyperparameters

The lack of ground truth for a real-world dataset like cdMRI makes quantitative analysis difficult. The qualitative comparison between subjects, scanners, and acquisitions is shown in Figure 9. We see comparable size and sharpness of FODs for both INR-CSD fits between subjects. Lower spatial resolution (in the standard acquisitions) also leads to decreased sharpness of the FODs. The voxel-wise CSD fit on subject K appears to have sharper FODs but displays spurious, less aligned peaks (green arrow in Figure 9) compared to the INR-CSD fit.

**Figure 9:**
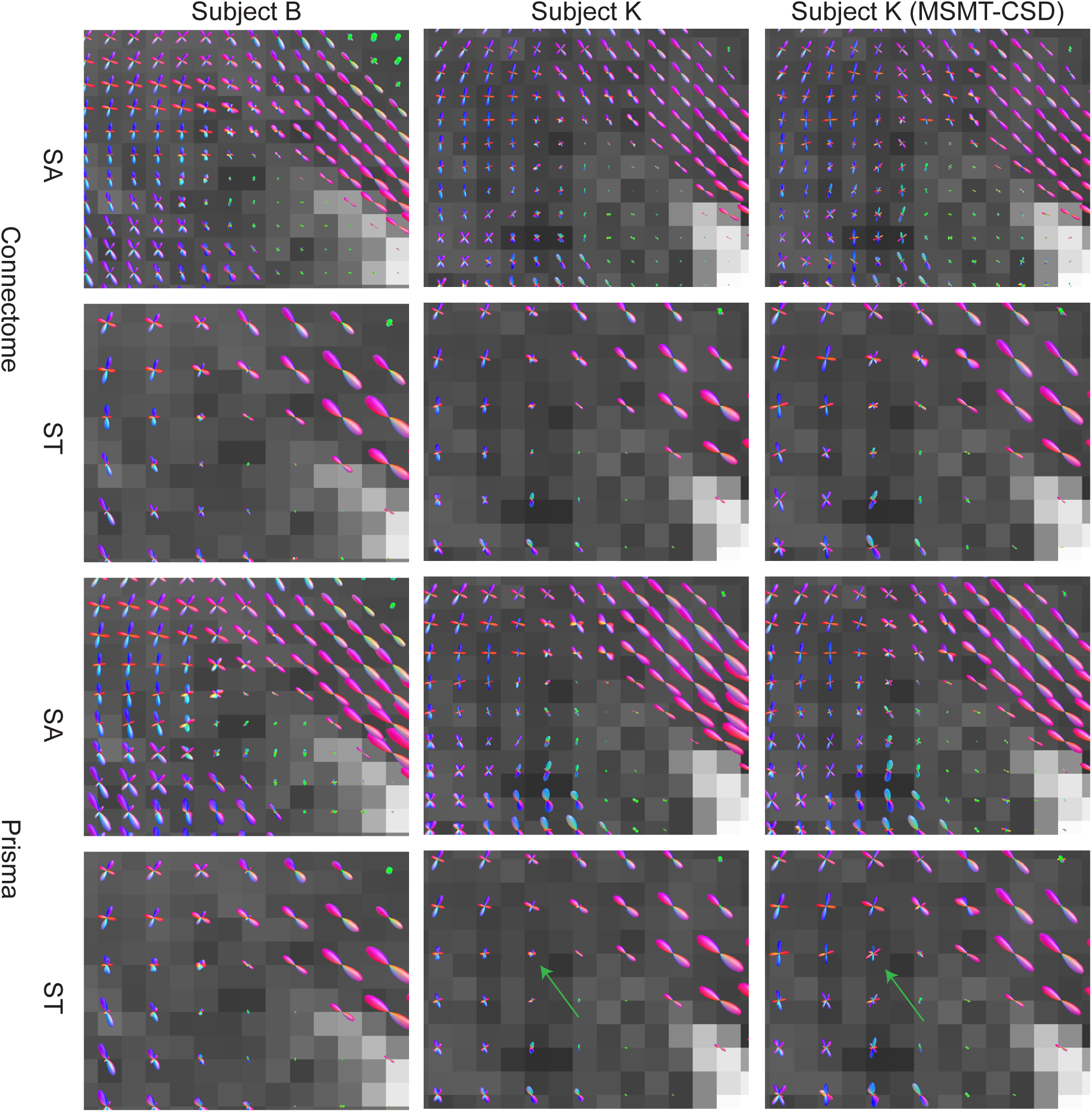
Visualizations of output FODs in the right periventricular area for the INR-MSMT model for different subjects (B and K), scanners (Connectom and Prisma), and acquisitions (state of the art (SA) and standard (ST)). Voxel-wise MSMT-CSD serves as a comparison for one of the subjects. The background shows the T1 weighted image. The green arrow shows an example of an FOD that is less sharp in the INR fit but displays less spurious peaks.

### Real-world application of the model

The FODs for INR-CSD, linearly interpolated CSD, and MSMT-CSD are shown in Figure 10. In the CSO (A), we see fewer spurious peaks in the INR fit than in the voxel-wise CSD, especially in the ascending/descending corticospinal tract. Compared to MSMT-CSD, the INR and voxel-wise CSD show less variability between voxels. In area B, the INR fit has a sharper border between the corpus callosum and the CSF in the ventricle than the voxel-wise fit, as indicated by the yellow line in Figure 10B. Area C again shows a reduction in spurious peaks from the INR-fit to the voxel-wise fit, best seen below the apex of the sulcus (yellow box in Figure 10C). Compared to MSMT-CSD, there is increased coherence between voxels in the INR-CSD but decreased FOD sharpness.

**Figure 10:**
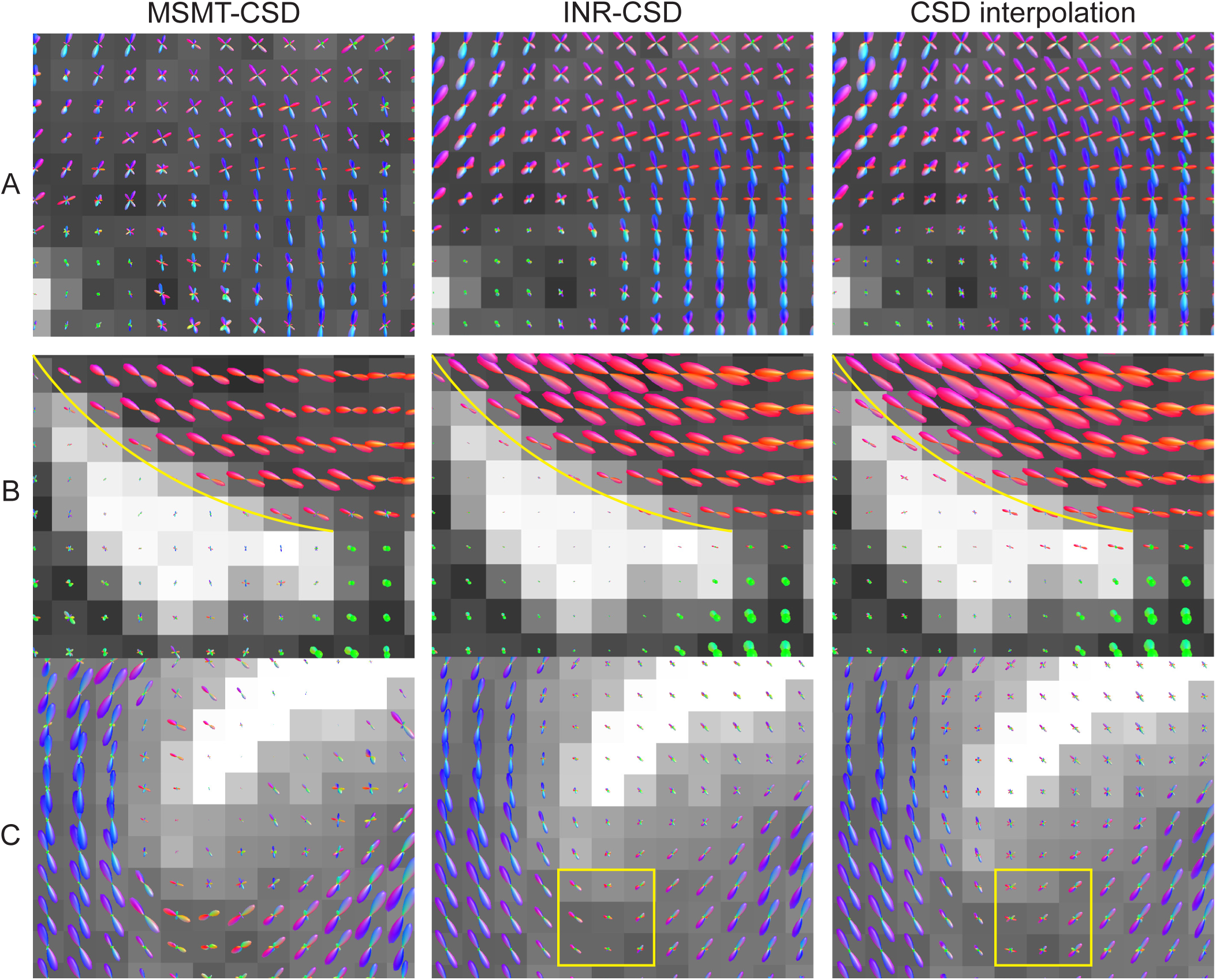
Visualizations of the output FODs in the centrum semi-ovale (A), the peri-ventricular bundles (B) and around a sulcus (C), for multi-shell multi-tissue CSD (MSMT-CSD) on three shells of the 1.2mm Connectom dataset, and INR-CSD and voxel-wise CSD fit on a single *b* = 3000*m/s*^2^ shell of the 2.4mm Connectom dataset. The background shows the T1 weighted image. The yellow line shows the border between the corpus callosum and the ventricle. The yellow box shows a sharply curved region where INR-CSD creates a smoother curve compared to voxel-wise CSD.

## 5 Discussion

### Implications of results

In this work, we propose INR-CSD and INR-MSMT, an approach to create a continuous representation of FODs from a dMRI dataset by combining (MSMT-)CSD with INRs. Our approach differs from recently proposed INR approaches (Consagra et al., 2024; Hendriks et al., 2023). Most importantly, we apply a model-based approach, which allows us to apply dataset and outcome-specific priors (e.g., the response functions) to the training process using multiple shells. The INR outputs the SH-coefficients of the FOD at the desired location directly, bypassing the need for interpolation. The INR can produce FODs representing the underlying structures, even in noisy environments, and when sampled at resolutions significantly higher than the input. Fitting the model to the data can be done in seconds to minutes on consumer-grade hardware based on the input resolution and the number of gradient directions. This, alongside the displayed robustness of the hyperparameter settings, shows that our method can be feasibly used in daily practice. Our method produces similar or better quantitative results in all experiments on synthetic datasets for which the ground truth is available. The visualizations show improvement compared to the baseline in the spatial interpolation of highly curved fibers, where fewer spurious peaks are produced. Noisy (real-world) acquisitions are modeled more consistently in our approach, showing more coherence between voxels and sharper interfaces between bundles and tissues. These results indicate that the proposed method increases the abilities of the model to estimate the coefficients of the FOD based on the underlying structure. The differences between our suggested method and the baseline are the smallest when dealing with angular sparsity: areas with low crossing angles produce sharper FODs using the baseline method, but areas of high curvature produce more spurious peaks. In general, the FODs produced by the INR appear slightly less sharp but with more intervoxel coherence than the FODs from CSD.

### Limitations of the approach

The methods in this paper are based on early INR literature (Mildenhall et al., 2020; Tancik et al., 2020), after which many alternatives have been suggested (Molaei et al., 2023). The positional encoding, model architecture, and training process can all be subject to improvements. An example of a possible improvement would be the sinusoidal networks used by Consagra et al., 2024 and Mancini et al., 2022. We should not only consider applying the existing state-of-the-art but also look into dMRI-specific improvements (e.g., positional encodings based on bundle shapes or dMRI-specific loss functions). Additionally, the model outputs FODs directly, which limits the usability of this approach to scenarios where downstream FOD metrics or fiber tracking is the end goal. In other scenarios, we can revert to the models introduced in previous work that output dMRI signal or use an INR to parameterize different dMRI models (Alexander et al., 2019).

While fiber tracking is the most obvious use case for our proposed models, we have chosen not to include actual fiber tracking experiments in this work. We believe that tracking on a continuous space requires a novel set of tracking methodologies - that goes beyond traditional step-wise streamline methods. The evaluation of the results would also be challenging: many comparisons to existing methods can be made (i.e., different fiber tracking algorithms and different interpolation methods), for which the computational speed and accuracy depend on the implementation. Nonetheless, we have performed a preliminary analysis of fiber tracking using our methods, which is discussed in Appendix B.

We designed the synthetic experiments to separate the different difficulties a model can encounter while fitting a dataset. This is both a strength and a weakness. It allows for quantitative analysis of the model and its comparison in a controlled environment. However, in a real dataset, low spatial and angular resolution and noise artifacts do not present themselves separately, nor are the issues limited to these three. The real data experiments overcome this limitation somewhat but do not allow us to analyze the results quantitatively. Whether the FODs obtained through MSMT-CSD in these experiments are the ground truth is debatable since imperfections in response function and partial volume estimations can lead to inaccuracies. We can see a higher estimation of the GM fraction by the INR compared to MSMT-CSD. This could be an area of improvement for the INR fit if we consider MSMT-CSD to be the ground truth for the volume fractions. Possibly, tuning the hyperparameters of the fitting process or changing the model architecture can generate a similar output to MSMT-CSD. If, however, the true volume fraction indeed lies higher, fitting using an INR (in this case) would be an improvement over the existing method. Furthermore, during one of the synthetic experiments, we applied CSD to a noisy dataset, while it would be common to first denoise the data before doing so (Veraart et al., 2016). This is done to showcase the denoising abilities of the INR and make a comparison between methods when fit on ‘raw’ data. Finally, although all three quantitative metrics are commonly used, they do not provide a perfect comparison between FODs. For example, NuFO can vary considerably based on the threshold values for a peak. This is seen in the angular sparsity experiment, where minor differences in FODs (small peaks or slightly different lobe orientations) change the NuFO from 1 to 2 or vice versa.

### Future directions

Everything considered, the usage of CSD model-based INRs for dMRI data is feasible, and it shows great promise for handling spatial sparsity and noise in particular. There are many highly effective upsampling and denoising techniques available (Alexander et al., 2017; Dyrby et al., 2014; Tax et al., 2022), but using INRs has specific advantages compared to these methods. For one, the output of the INR can be tailored to the desired diffusion model output, i.e., after fitting once, the inference immediately produces the requested outcome. Future work would be necessary, comparing the quality and speed of computation for existing interpolation and denoising methods and the inference quality and speed of an INR model, which are highly dependent on the encoding and model sizes. Secondly, the ability to sample in a continuous space is a significant advantage. One can imagine probabilistic fiber tracking using our models, benefiting from the computational efficiency and high-quality representation of the INRs. Thirdly, the INR and spatial encoding provide the user with hyperparameters that can be fine-tuned to fit their specific needs. We have analyzed the effects of different hyperparameters in Appendix A, but further research might be necessary to pinpoint exactly how they can influence the results. Some examples could be using lower frequency encodings (by lowering sigma) if we are only interested in the more extensive tracts or coarse structures. Or decreasing the model or encoding sizes to create lower complexity representations in scenarios with poor-quality datasets. Finally, and most interesting, is the possibility, as suggested by Consagra et al., 2024, that the spatial encoding allows the model to learn spatial correlations in the dataset. The results in the spatial resolution, noise, and cdMRI experiment especially point toward the INR having a notion of structural coherence it can exploit to create a better representation. In our model specifically, we see that the INR can capture curved fibers accurately at a resolution much higher than the input. Curvature, fanning, crossing, and kissing fibers are difficult to deal with at a voxel level. This has resulted in the suggestion of using asymmetric FODs, which make use of neighborhood information to more accurately represent the fiber populations in the voxel (Barmpoutis et al., 2008; Bastiani et al., 2017; Poirier & Descoteaux, 2024; Reisert et al., 2012; Wu et al., 2020). Theoretically, a well-fit INR that correctly captures the underlying structure could circumvent the usage of asymmetrical FODs by sampling the regions at a sufficiently high resolution. More interestingly, we could apply INRs to the work done on asymmetrical FODs to further improve the quality of the INR. The improved representation through spatial correlations is not limited to INR-CSD and INR-MSMT, however, but could be used to fit the parameters of any dMRI model. This could increase fit quality, especially when fitting more data-demanding models (e.g., the standard model, Novikov et al., 2019).

## 6 Conclusion

This work introduces the INR-CSD and MSMT-CSD models, which integrate the models of CSD and MSMT-CSD with recent research on INRs in dMRI, highlighting both their promising capabilities and limitations. Notably, the denoising abilities and precise *on the fly* interpolation of FODs offer valuable starting points for further investigation. Ultimately, this research paves the way for applying INRs to other dMRI models, which may benefit from the spatially informed parameter estimation provided by an INR approach compared to a voxel-wise method.

## Data and Code Availability

The ISMRM dataset was obtained from: https://tractometer.org/ismrm2015/home/. The CDMRI dataset was obtained from: https://www.cardiff.ac.uk/cardiff-university-brain-research-imaging-centre/research/projects/cross-scanner-and-cross-protocol-diffusion-MRI-data-harmonisation. The code will be made available to the community via GitHub upon publication at: https://github.com/tomhend/MSMT-CSD INR.

## Ethics statement

This work was conducted per approval from the Eindhoven University of Technology Ethical Review Board (ERB2023MCS5).

## Author Contributions

TH conceived and implemented the method; performed the experiments; analyzed the results, created the figures, and wrote the first draft of the manuscript. AV and MC provided guidance in data use and assisted in experimental design and results interpretation. All authors contributed to revisions of the manuscript prior to submission.

## Acknowledgements

The CDMRI data were acquired at the UK National Facility for In Vivo MR Imaging of Human Tissue Microstructure located in CUBRIC funded by the EPSRC (grant EP/M029778/1), and The Wolfson Foundation. Acquisition and processing of the data was supported by a Rubicon grant from the NWO (680-50-1527), a Wellcome Trust Investigator Award (096646/Z/11/Z), and a Wellcome Trust Strategic Award (104943/Z/14/Z). This database was initiated by the 2017 and 2018 MICCAI Computational Diffusion MRI committees (Chantal Tax, Francesco Grussu, Enrico Kaden, Lipeng Ning, Jelle Veraart, Elisenda Bonet-Carne, and Farshid Sepehrband) and CUBRIC, Cardiff University (Chantal Tax, Derek Jones, Umesh Rudrapatna, John Evans, Greg Parker, Slawomir Kusmia, Cyril Charron, and David Linden). We also thank Ruben Vink for his input and interesting discussions.

## A Effect of hyperparameters

**Figure 11:**
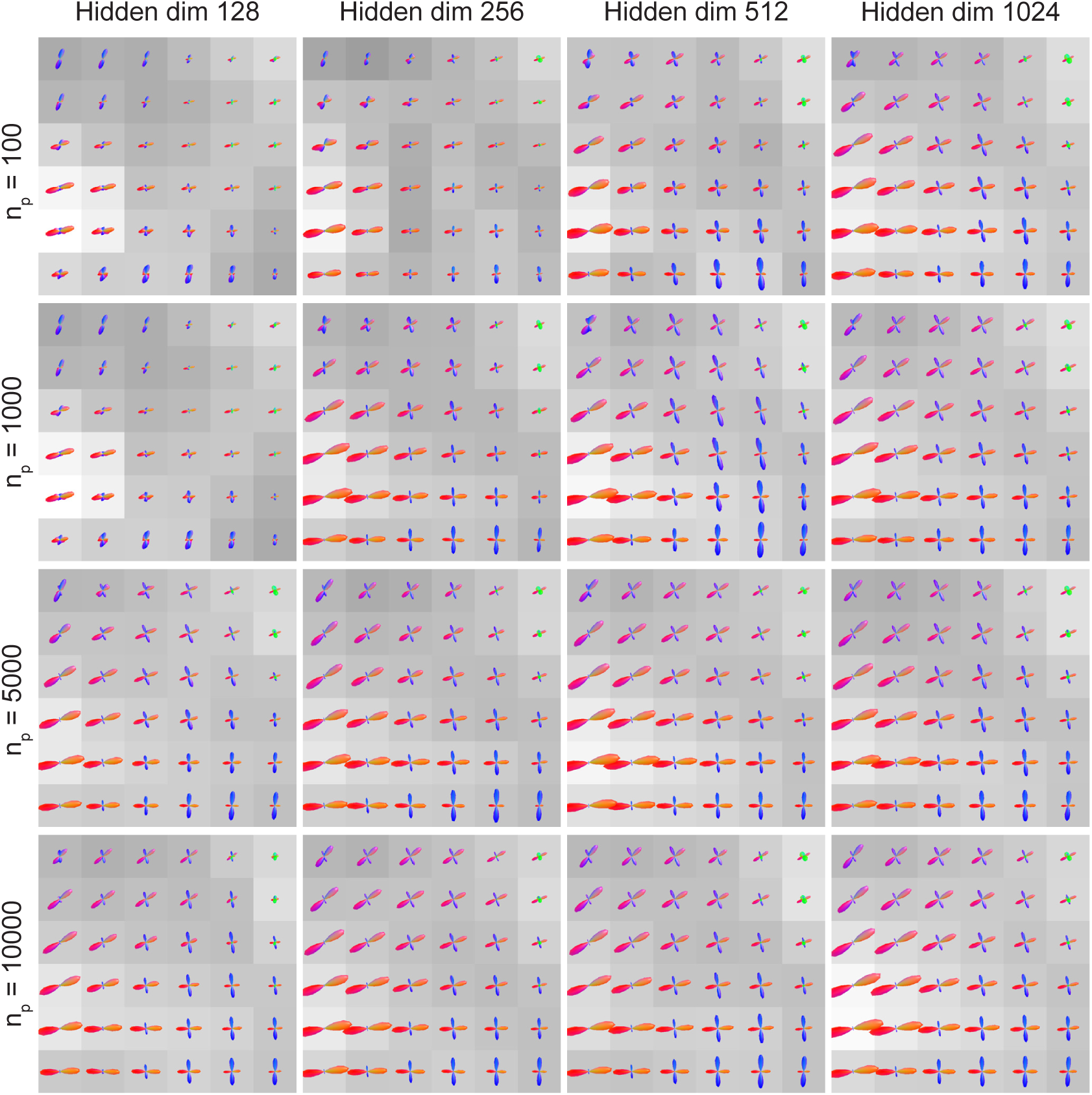
Visualizations of the FODs in the center semi-ovale for different combinations of *n_p_* (rows) and hidden dimensions (columns), at a constant sigma of 4. Increasing either parameter leads to an improvement in FOD quality. Increasing *n_p_* has a smaller effect on model size and fitting time and is preferable over increasing the number of hidden dimensions.

**Figure 12:**
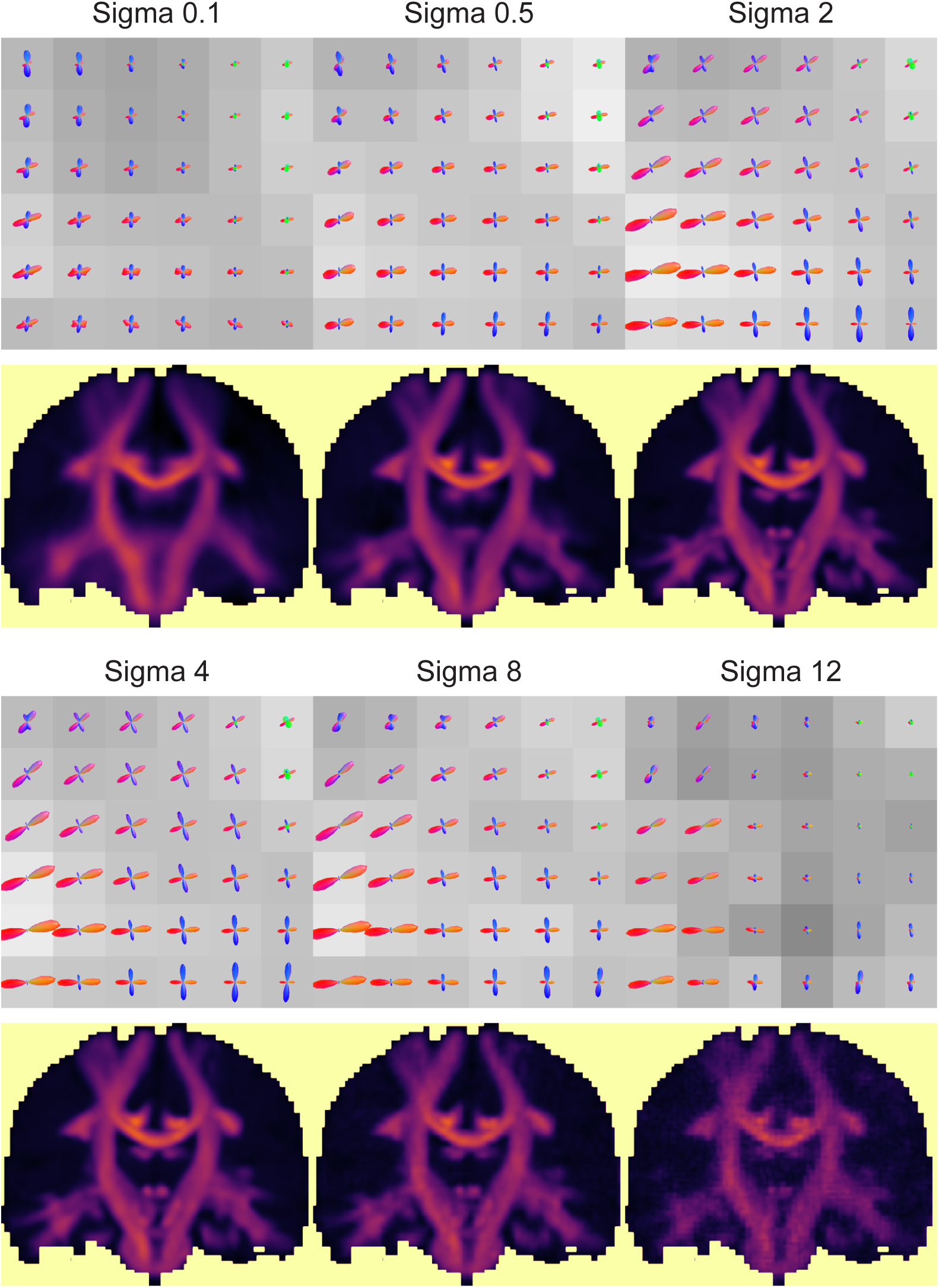
Visualizations of FODs in the center semi-ovale, as well as an overview of the AFD map, for changing values for sigma, at a constant 1024 hidden dimensions and 5000 positional encodings. Lower sigma values show a high similarity between voxels and a blurred AFD map. In contrast, high sigma values show a low similarity between voxels and a more granular AFD map. Sigma = 2 and sigma = 4 show a nice balance between intervoxel coherence and smoothness of the AFD map.

## B Preliminary fiber tracking results

This appendix provides some preliminary results and quantitative measures concerning fiber tracking for our proposed models. The initial fit of the INR on a considerably high resolution (1.25mm isotropic) dataset with an average number of gradient directions (30 or 60 non *b* = 0*s/mm*^2^ images) was acquired in 3 to 5.5 minutes for INR-CSD and INR-MSMT respectively. This is considerably faster than previous work using INRs for diffusion MRI (Consagra et al., 2024). After the initial fit, a batch of 100,000 FODs can be produced at arbitrary locations in the volume in 0.8ms once the data is on the GPU using the Pytorch library. Moving 100,000 coordinates from the CPU to the GPU, running the model for the coordinates, and moving back the FODs from the GPU to the CPU increases this time to 150ms.

While a complete evaluation of tracking in the continuous space is too elaborate and out of scope for this work, we want to provide insight into the potential qualitative improvements with a simple experiment. We fit the INR-CSD and voxel-wise CSD on the 2.5mm resolution noiseless synthetic dataset using 30 non *b* = 0*s/mm*^2^ directions. The INR-CSD is then sampled at 1.25mm resolution, and the CSD FODs are upsampled to 1.25mm resolution using linear interpolation. Both FODs are then input into the MRtrix implementation of the iFOD2 algorithm to perform full brain fiber tracking, using GM-WM interface seeding regions, stopping when 2M valid streamlines are reached (J. D. Tournier et al., 2010). This implementation uses linear interpolation between the FODs. This implies that any difference in output (excluding the stochasticity inherent to the iFOD2 algorithm) is because of differences in the upsampling step from 2.5mm to 1.25mm isotropic resolution. We compare the output to the ground-truth streamlines used to generate the synthetic dataset. The results can be seen in Figure 13. We can see that both INR-CSD and CSD show partial voluming of the fiber bundles into regions that have no bundles in the ground truth. In both regions, this appears to be less profound in the INR-CSD than in the CSD fiber tracking. Possibly, these differences can be more significant when tracking in the continuous space, as we are still relying on linear interpolation for the sub 1.25mm FODs.

Finally, we use the Tractometer created for the ISMRM 2015 challenge to score both tractograms (Maier-Hein et al., 2017; Renauld et al., 2023). Metrics include the number of valid bundles found (VB, out of the 21 ground truth bundles), the number of valid and invalid streamlines (VS, IS), the overlap (OL, the percentage of voxels covered by the ground truth which are also covered by the tractogram), the normalized overreach (ORn, the percentage of voxels covered outside of the ground truth, normalized by the ground truth size), and the F1-score. An in-depth explanation of these metrics can be found in Renauld et al., 2023. Table 2 shows the INR-CSD and the voxel-wise CSD approach results. CSD finds more valid streamlines but produces the same OL and a higher ORn. This confirms the findings of the qualitative comparison, as these scores indicate that more streamlines are going outside the ground truth bundles. The resulting F1-score is, therefore, higher in INR-CSD compared to CSD.

**Figure 13:**
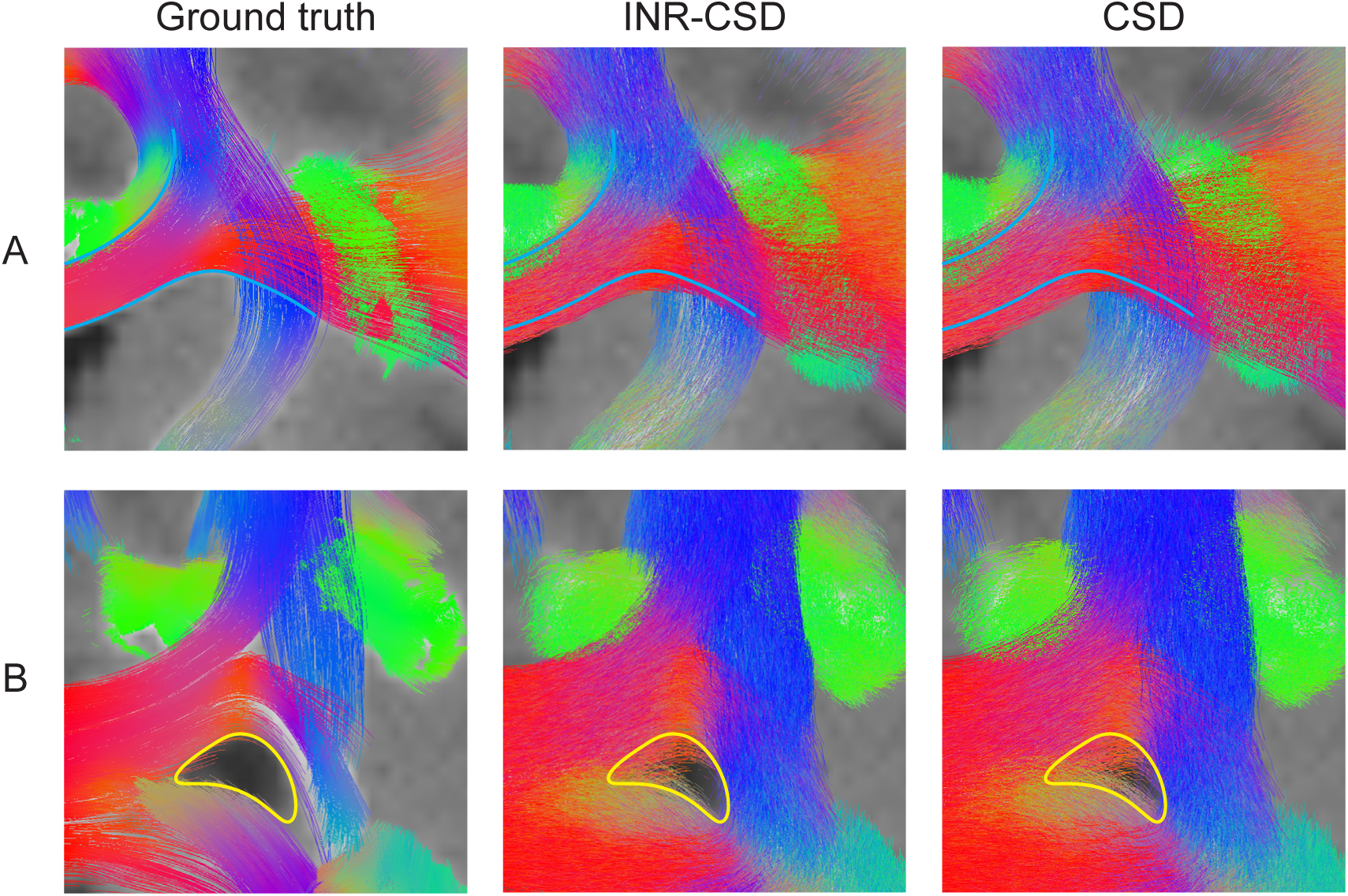
Visualizations of streamlines of the ground truth fibers, the INR-CSD tracking, and the CSD tracking for the centrum semi-ovale (A) and a periventricular region (B).

**Table 2:**
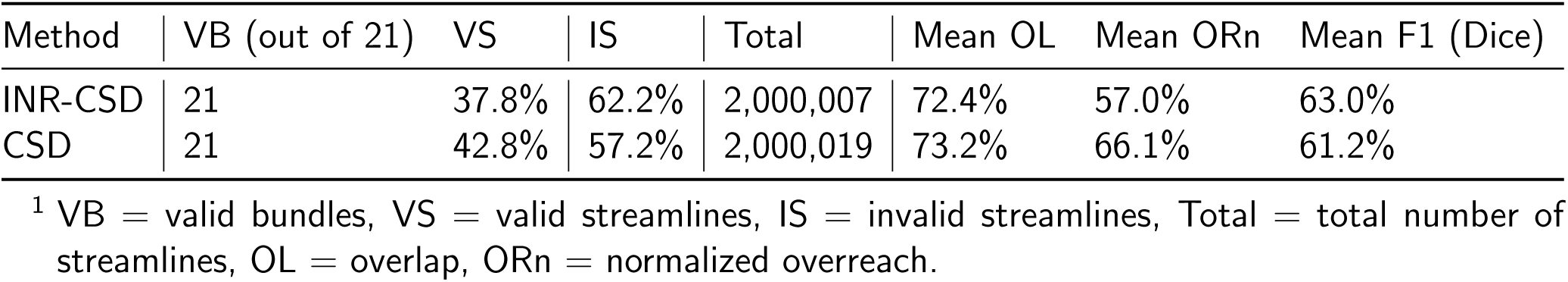
Tractometer scores^1^.

| Method | VB (out of 21) | VS | IS | Total | Mean OL | Mean ORn | Mean F1 (Dice) |
| --- | --- | --- | --- | --- | --- | --- | --- |
| INR-CSD | 21 | 37.8% | 62.2% | 2,000,007 | 72.4% | 57.0% | 63.0% |
| CSD | 21 | 42.8% | 57.2% | 2,000,019 | 73.2% | 66.1% | 61.2% |

<sup>1</sup> VB = valid bundles, VS = valid streamlines, IS = invalid streamlines, Total = total number of streamlines, OL = overlap, ORn = normalized overreach.

## Footnotes

Note: different definitions for *l* and *m* are used throughout literature. We adopt the definitions used by MRtrix3 (J.-D. Tournier et al., 2019).

https://github.com/scipy/scipy version 1.9.3

https://github.com/pytorch/pytorch version 2.0.0

https://github.com/scilus/scilpy version 2.0.1

## References

Alexander, D. C., Dyrby, T. B., Nilsson, M., & Zhang, H. (2019). Imaging brain microstructure with diffusion mri: Practicality and applications. NMR in Biomedicine, 32 (4), e3841.

Alexander, D. C., Zikic, D., Ghosh, A., Tanno, R., Wottschel, V., Zhang, J., Kaden, E., Dyrby, T. B., Sotiropoulos, S. N., Zhang, H., et al. (2017). Image quality transfer and applications in diffusion mri. NeuroImage, 152, 283–298.

Anderson, A. W. (2005). Measurement of fiber orientation distributions using high angular resolution diffusion imaging. Magnetic Resonance in Medicine, 54 (5), 1194–1206. Retrieved July 22, 2024, from https://onlinelibrary.wiley.com/doi/abs/10.1002/mrm.20667

Barmpoutis, A., Vemuri, B. C., Howland, D., & Forder, J. R. (2008). Extracting tractosemas from a displacement probability field for tractography in dw-mri. International Conference on Medical Image Computing and Computer-Assisted Intervention, 9–16.

Bastiani, M., Cottaar, M., Dikranian, K., Ghosh, A., Zhang, H., Alexander, D. C., Behrens, T. E., Jbabdi, S., & Sotiropoulos, S. N. (2017). Improved tractography using asymmetric fibre orientation distributions. Neuroimage, 158, 205–218.

Consagra, W., Ning, L., & Rathi, Y. (2024). Neural orientation distribution fields for estimation and uncertainty quantification in diffusion MRI. Medical Image Analysis, 93, 103105. Retrieved July 22, 2024, from https://www.sciencedirect.com/science/article/pii/S1361841524000306

Dell’Acqua, F., Simmons, A., Williams, S. C., & Catani, M. (2013). Can spherical deconvolution provide more information than fiber orientations? Hindrance modulated orientational anisotropy, a true-tract specific index to characterize white matter diffusion. Human Brain Mapping, 34 (10), 2464– 2483. Retrieved July 22, 2024, from https://onlinelibrary.wiley.com/doi/abs/10.1002/hbm.22080

Dhollander, T., Mito, R., Raffelt, D., & Connelly, A. (2019). Improved white matter response function estimation for 3-tissue constrained spherical deconvolution.

Dyrby, T. B., Lundell, H., Burke, M. W., Reislev, N. L., Paulson, O. B., Ptito, M., & Siebner, H. R. (2014). Interpolation of diffusion weighted imaging datasets. Neuroimage, 103, 202–213.

Elaldi, A., Dey, N., Kim, H., & Gerig, G. (2021). Equivariant Spherical Deconvolution: Learning Sparse Orientation Distribution Functions from Spherical Data. In A. Feragen, S. Sommer, J. Schnabel, & M. Nielsen (Eds.), Information Processing in Medical Imaging (pp. 267–278). Springer International Publishing. 10.1007/978-3-030-78191-021

Elaldi, A., Gerig, G., & Dey, N. (2024). E(3) × SO(3)-Equivariant Networks for Spherical Deconvolution in Diffusion MRI. Proceedings of machine learning research, 227, 301–319. Retrieved July 22, 2024, from https://www.ncbi.nlm.nih.gov/pmc/articles/PMC10901527/

Genc, S., Tax, C. M., Raven, E. P., Chamberland, M., Parker, G. D., & Jones, D. K. (2020). Impact of b-value on estimates of apparent fibre density. Human brain mapping, 41 (10), 2583–2595.

Hendriks, T., Vilanova, A., & Chamberland, M. (2023). Neural Spherical Harmonics for Structurally Coherent Continuous Representation of Diffusion MRI Signal. In M. Karaman, R. Mito, E. Powell, F. Rheault, & S. Winzeck (Eds.), Computational Diffusion MRI (pp. 1–12). Springer Nature Switzerland. 10.1007/978-3-031-47292-31

Jeurissen, B., Descoteaux, M., Mori, S., & Leemans, A. (2019). Diffusion mri fiber tractography of the brain. NMR in Biomedicine, 32 (4), e3785.

Jeurissen, B., Leemans, A., Tournier, J.-D., Jones, D. K., & Sijbers, J. (2013). Investigating the prevalence of complex fiber configurations in white matter tissue with diffusion magnetic resonance imaging. Human Brain Mapping, 34 (11), 2747–2766. Retrieved July 22, 2024, from https://onlinelibrary.wiley.com/doi/abs/10.1002/hbm.22099

Jeurissen, B., Tournier, J.-D., Dhollander, T., Connelly, A., & Sijbers, J. (2014). Multi-tissue constrained spherical deconvolution for improved analysis of multi-shell diffusion MRI data. NeuroImage, 103, 411–426. Retrieved July 22, 2024, from https://www.sciencedirect.com/science/article/pii/S1053811914006442

Karimi, D. (2024, January 1). Diffusion MRI with Machine Learning. arXiv: 2402.00019 [cs, eess]. Retrieved July 22, 2024, from http://arxiv.org/abs/2402.00019

Maier-Hein, K. H., Neher, P. F., Houde, J.-C., Côté, M.-A., Garyfallidis, E., Zhong, J., Chamberland, M., Yeh, F.-C., Lin, Y.-C., Ji, Q., et al. (2017). The challenge of mapping the human connectome based on diffusion tractography. Nature communications, 8 (1), 1349.

Mancini, M., Jones, D. K., & Palombo, M. (2022). Lossy Compression of Multidimensional Medical Images Using Sinusoidal Activation Networks: An Evaluation Study. In S. Cetin-Karayumak, D. Christiaens, M. Figini, P. Guevara, T. Pieciak, E. Powell, & F. Rheault (Eds.), Computational Diffusion MRI (pp. 26–37). Springer Nature Switzerland. 10.1007/978-3-031-21206-23

Mildenhall, B., Srinivasan, P. P., Tancik, M., Barron, J. T., Ramamoorthi, R., & Ng, R. (2020, August 3). NeRF: Representing Scenes as Neural Radiance Fields for View Synthesis. arXiv: 2003.08934 [cs]. Retrieved May 11, 2023, from http://arxiv.org/abs/2003.08934

Molaei, A., Aminimehr, A., Tavakoli, A., Kazerouni, A., Azad, B., Azad, R., & Merhof, D. (2023). Implicit neural representation in medical imaging: A comparative survey. Proceedings of the IEEE/CVF International Conference on Computer Vision (ICCV) Workshops, 2381–2391.

Neher, P. F., Laun, F. B., Stieltjes, B., & Maier-Hein, K. H. (2014). Fiberfox: Facilitating the creation of realistic white matter software phantoms. Magnetic Resonance in Medicine, 72 (5), 1460–1470. 10.1002/mrm.25045

Nie, X., & Shi, Y. (2023). Flow-Based Geometric Interpolation of Fiber Orientation Distribution Functions. In H. Greenspan, A. Madabhushi, P. Mousavi, S. Salcudean, J. Duncan, T. Syeda-Mahmood, & R. Taylor (Eds.), Medical Image Computing and Computer Assisted Intervention – MICCAI 2023 (pp. 46–55). Springer Nature Switzerland. 10.1007/978-3-031-43993-35

Ning, L., Bonet-Carne, E., Grussu, F., Sepehrband, F., Kaden, E., Veraart, J., Blumberg, S. B., Khoo, C. S., Palombo, M., Kokkinos, I., et al. (2020). Cross-scanner and cross-protocol multi-shell diffusion mri data harmonization: Algorithms and results. Neuroimage, 221, 117128.

Novikov, D. S., Fieremans, E., Jespersen, S. N., & Kiselev, V. G. (2019). Quantifying brain microstructure with diffusion MRI: Theory and parameter estimation. NMR in Biomedicine, 32 (4), e3998. Retrieved December 18, 2023, from https://onlinelibrary.wiley.com/doi/abs/10.1002/nbm.3998

Poirier, C., & Descoteaux, M. (2024). A unified filtering method for estimating asymmetric orientation distribution functions. NeuroImage, 287, 120516.

Raffelt, D., Tournier, J.-., Rose, S., Ridgway, G. R., Henderson, R., Crozier, S., Salvado, O., & Connelly, A. (2012). Apparent Fibre Density: A novel measure for the analysis of diffusion-weighted magnetic resonance images. NeuroImage, 59 (4), 3976–3994. Retrieved July 22, 2024, from https://www.sciencedirect.com/science/article/pii/S1053811911012092

Reisert, M., Kellner, E., & Kiselev, V. G. (2012). About the geometry of asymmetric fiber orientation distributions. IEEE transactions on medical imaging, 31 (6), 1240–1249.

Renauld, E., Théberge, A., Petit, L., Houde, J.-C., & Descoteaux, M. (2023). Validate your white matter tractography algorithms with a reappraised ISMRM 2015 Tractography Challenge scoring system. Scientific Reports, 13 (1), 2347. Retrieved July 22, 2024, from https://www.nature.com/articles/s41598-023-28560-w

Schilling, K. G., Gao, Y., Li, M., Wu, T.-L., Blaber, J., Landman, B. A., Anderson, A. W., Ding, Z., & Gore, J. C. (2019). Functional tractography of white matter by high angular resolution functional-correlation imaging (harfi). Magnetic resonance in medicine, 81 (3), 2011–2024.

Schilling, K. G., Nath, V., Blaber, J., Harrigan, R. L., Ding, Z., Anderson, A. W., & Landman, B. A. (2017). Effects of b-value and number of gradient directions on diffusion mri measures obtained with q-ball imaging. Medical Imaging 2017: Image Processing, 10133, 179–185.

Tancik, M., Srinivasan, P. P., Mildenhall, B., Fridovich-Keil, S., Raghavan, N., Singhal, U., Ramamoorthi, R., Barron, J. T., & Ng, R. (2020, June 18). Fourier Features Let Networks Learn High Frequency Functions in Low Dimensional Domains. arXiv: 2006.10739 [cs]. Retrieved May 15, 2023, from http://arxiv.org/abs/2006.10739

Tax, C. M., Bastiani, M., Veraart, J., Garyfallidis, E., & Irfanoglu, M. O. (2022). What’s new and what’s next in diffusion mri preprocessing. NeuroImage, 249, 118830.

Tax, C. M., Grussu, F., Kaden, E., Ning, L., Rudrapatna, U., John Evans, C., St-Jean, S., Leemans, A., Koppers, S., Merhof, D., Ghosh, A., Tanno, R., Alexander, D. C., Zappalà, S., Charron, C., Kusmia, S., Linden, D. E., Jones, D. K., & Veraart, J. (2019). Cross-scanner and cross-protocol diffusion MRI data harmonisation: A benchmark database and evaluation of algorithms. NeuroImage, 195, 285–299. Retrieved July 22, 2024, from https://www.sciencedirect.com/science/article/pii/S1053811919300837

Tournier, J. D., Calamante, F., Connelly, A., et al. (2010). Improved probabilistic streamlines tractography by 2nd order integration over fibre orientation distributions. Proceedings of the international society for magnetic resonance in medicine, 1670.

Tournier, J.-D., Calamante, F., & Connelly, A. (2007). Robust determination of the fibre orientation distribution in diffusion MRI: Non-negativity constrained super-resolved spherical deconvolution. NeuroImage, 35 (4), 1459–1472. Retrieved July 22, 2024, from https://www.sciencedirect.com/science/article/pii/S1053811907001243

Tournier, J.-D., Calamante, F., & Connelly, A. (2013). Determination of the appropriate b value and number of gradient directions for high-angular-resolution diffusion-weighted imaging. NMR in Biomedicine, 26 (12), 1775–1786. Retrieved July 22, 2024, from https://onlinelibrary.wiley.com/doi/abs/10.1002/nbm.3017

Tournier, J.-D., Smith, R., Raffelt, D., Tabbara, R., Dhollander, T., Pietsch, M., Christiaens, D., Jeurissen, B., Yeh, C.-H., & Connelly, A. (2019). MRtrix3: A fast, flexible and open software framework for medical image processing and visualisation. NeuroImage, 202, 116137. 10.1016/j.neuroimage.2019.116137

Veraart, J., Novikov, D. S., Christiaens, D., Ades-Aron, B., Sijbers, J., & Fieremans, E. (2016). Denoising of diffusion mri using random matrix theory. Neuroimage, 142, 394–406.

Wu, Y., Hong, Y., Feng, Y., Shen, D., & Yap, P.-T. (2020). Mitigating gyral bias in cortical tractography via asymmetric fiber orientation distributions. Medical image analysis, 59, 101543.

